# HIF1α controls somitogenesis and spine development by regulating levels of intracellular oxygen in the presomitic mesoderm

**DOI:** 10.64898/2025.12.19.694009

**Authors:** Matthew J. Anderson, Angela Yao, Brittany Laslow, Ernestina Schipani, Mark Lewandoski

## Abstract

Embryonic axis extension depends on presomitic mesoderm (PSM) segmentation into somites, which give rise to vertebrae and muscles. The PSM is a highly glycolytic structure. HIF1α is a transcription factor that regulates bioenergetic metabolism by promoting glycolysis and suppressing mitochondrial respiration. Conditional deletion of HIF1α in the PSM resulted in vertebral malformations, including shortened, misshapen vertebrae and additional thoracic segments. These defects correlated with disrupted segmentation clock oscillations, marked by abnormal *Hes7* and *Lfng* expression. HIF1α loss also induced intracellular hypoxia, likely due to increased mitochondrial respiration. Hyperoxic exposure of mutant embryos corrected hypoxia, restoring *Hes7* oscillations and somitogenesis. These findings highlight HIF1α’s essential role in ensuring proper segmentation clock function and somitogenesis by maintaining intracellular oxygen homeostasis through reprogramming of metabolism.

## Introduction

The formation of the vertebrate axis integrates numerous pathways and processes, with mesoderm formation and subsequent somite development being among the earliest steps. Somites are transient, segmental structures unique to chordates that give rise to critical components of the torso, including muscles, tendons, vertebrae, associated ribs, and the dermis [1]. During somitogenesis, somites are sequentially segmented from the presomitic mesoderm (PSM) at the embryo’s posterior. Disruptions in this process result in defects in the derived structures, particularly the vertebral column, and may lead to debilitating conditions such as congenital scoliosis [1, 2].

In mice, approximately 64 somites are formed over six days. To ensure all somites are formed, the PSM must be replenished in balance with somite segmentation. While these processes are interrelated, they are also distinct. Some mouse mutants exhibit defects in somite segmentation without impairments in mesoderm formation [3-5], whereas others show depletion of mesoderm progenitors with minimal effects on segmentation until the mesoderm is exhausted [6, 7]. However, most defects in mesoderm formation do impact somite segmentation due to the interconnectedness of the underlying pathways [8-10].

The widely accepted “Clock and Wavefront model” explains the coordination of mesoderm formation and somite segmentation [11]. The Clock activity regulates the timing of somite segmentation and differentiation in the anterior PSM, while the Wavefront activity determines the spatial position of segmentation by blocking differentiation signals in the posterior PSM thereby setting the anterior position of new somite segmentation. Molecular evidence for the Clock points to members of the Notch signaling pathway [12]. Notch signaling propagates through the PSM from posterior to anterior, forming oscillatory bands of activation and repression. When oscillations of activated Notch reach the anterior boundary of the Wavefront, they trigger the expression of segmentation genes like *Mesp2*, forming new somite boundaries[13, 14].

The Wavefront activity, primarily mediated by FGF and Wnt signaling, prevents premature differentiation in the PSM. Loss of *Fgf4* and *Fgf8* or deletion of *Wnt3a* or *β-Catenin* leads to premature differentiation, disrupting somitogenesis [8, 9]. Mesoderm levels are maintained during the first half of somitogenesis through gastrulation, which replenishes the PSM with new mesodermal cells as mesoderm is depleted in somitogenesis [15]. Gastrulation involves cells from the ectodermal epiblast layer undergoing epithelial-mesenchymal transition (EMT) and migrating into the underlying mesoderm layer. During this process, ectodermal genes such as *Sox2* and *E-Cadherin* are downregulated, while mesodermal genes, including *T (Brachyury)* and *Tbx6*, are upregulated [1, 16-18]. Disruptions in gastrulation lead to the premature cessation of somitogenesis and axis extension [18].

Recent studies have established a groundbreaking connection between cellular metabolism and somitogenesis [19, 20]. Cells in the posterior PSM primarily generate energy via glycolysis and lactate fermentation. Disruption of glycolysis or glucose deprivation leads to defective Notch oscillations [19, 21] and impaired mesoderm formation [20]. Glycolysis and lactate fermentation are critical processes in many progenitor cell types, including cancer cells, where metabolic intermediates are diverted from the TCA cycle for biosynthesis of lipids, proteins, and nucleotides [22] [23]. This phenomenon is called the Warburg effect. This process is modulated, in part, by the transcription factor HIF1α (Hypoxia Inducible Factor 1α) [24]. HIF1α enhances glycolytic enzyme expression, such as *Pgk1* and *Pkm* [25], while repressing oxidative phosphorylation [26]. Under normoxic conditions, HIF1α is ubiquitinated and degraded, but under hypoxia, it is stabilized [27].

Hypoxia has been shown to influence somitogenesis. Severe gestational hypoxia or mild hypoxia in genetically sensitized embryos negatively impacts segmentation clock oscillations [2]. Spondylocostal Dysostosis (SCDO) is characterized by severe vertebral malformations and is caused by homozygous loss-of-function (LOF) mutations of components of the Notch signaling pathway [28]. Mice carrying similar homozygous mutations phenocopy the human disease [2]. Heterozygous Notch LOF mutations cause the more modest, although more frequent, human defect of congenital scoliosis (CS), which is also phenocopied in heterozygous mouse mutants [28]. Those mouse phenotypes are worsened by gestational hypoxia [2]. Gestational hypoxia is a dangerous complication of pregnancy; it is due to a variety of pathological conditions including placental insufficiency and diabetes and can have profound long-term consequences [29].

Those findings suggest that HIF1α may be a critical regulator of metabolic and oxygen homeostasis during mesoderm formation and somitogenesis. *Hif1α* null homozygous embryos are not helpful in the testing of this hypothesis as they arrest at embryonic day 9.0 (E9.0) after forming only 11–12 somites and die within 48 hours [30].

To better understand the role of HIF1α in somitogenesis and mesoderm formation, as well as its involvement in metabolic regulation and hypoxia during these processes, we conditionally deleted *Hif1α* in the PSM. This approach aims to bypass early somitogenic failure and mid-gestational lethality, enabling a detailed investigation of HIF1α contributions to embryonic axis extension.

## Results

### Loss of HIF1α Results in Posterior PSM Hypoxia and Downregulation of Glycolytic Gene Expression

*Hif1α* is ubiquitously expressed throughout the embryo (Fig. S1), with higher levels observed in specific regions, such as the developing gut tube at E9.5 (Fig. S1C–D’) and the anterior presomitic mesoderm (PSM) at E10.5 (marked by an asterisk, Fig. S1F’). To investigate the role of HIF1α in somitogenesis and mesoderm formation, we conditionally deleted *Hif1α* in nascent mesoderm cells using TCre (Fig. S1G, H) [31]. Embryos with a TCre-mediated *Hif1α* deletion (TCre; *Hif1α* ^flox/null^) are referred to as “mutants”, and “controls” are littermate embryos (TCre; *Hif1α* ^flox/WT^) (Fig. S1G).

HIF1α is a key regulator of oxygen homeostasis, orchestrating vascular development and modulating metabolic gene expression. It achieves this by upregulating glycolytic pathway genes while repressing mitochondrial oxidative phosphorylation [25, 26, 30, 32]. In mutants, significant hypoxia is observed in the posterior mesoderm and neural tube at the 10–14 somite stage (ss) (Fig. 1A, B) and continues through ss 20-24 (Fig. C, D). Interestingly, tissues within the TCre lineage of control animals (carrying only one WT *Hif1a* allele) also exhibit slightly higher hypoxia compared to wild-type animals (Fig. 1C, E), whereas low-level hypoxia throughout the posterior WT embryo is only detectable with prolonged image acquisition (Fig. 1F). This suggests that HIF1α is specifically required to regulate intracellular oxygen levels in the posterior presomitic mesoderm (PSM) and neural tube.

**Fig 1.**
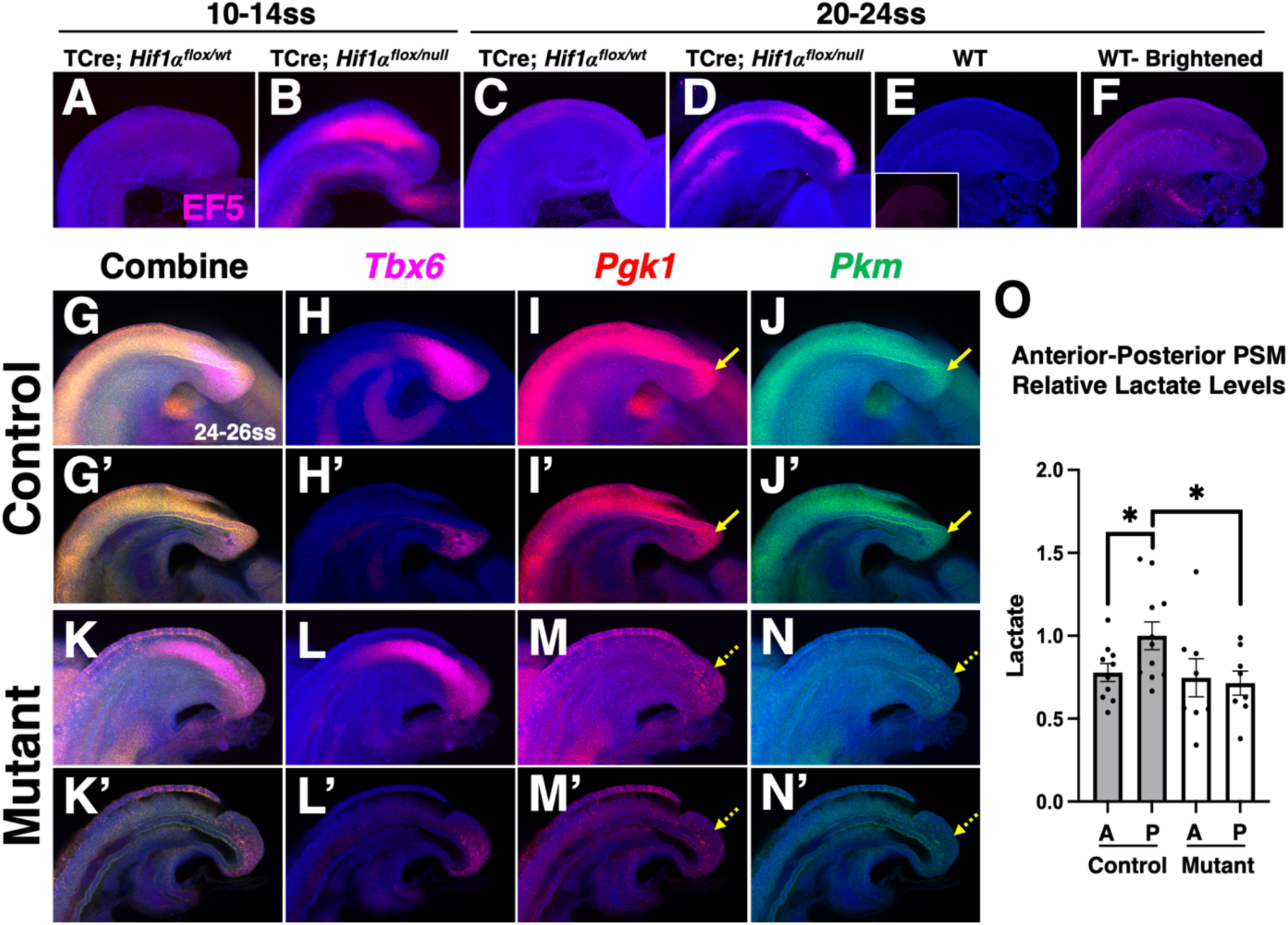
HIF1α mutants display increased hypoxia and reduced expression of glycolytic genes in the PSM. **(A-F)** EF5 staining for hypoxia in TCre; *Hif1α^fllox/wt^* (n= X in A; n= 5 in C), TCre; *Hif1α ^flox/null^* (N= X in B; n= 5 in D), or wildtype (n= 4) embryos at the somite stage indicated. D is same embryo in C with post-processing image brightening in to show low level hypoxia; lateral views of PSM, inset in E is EF5 staining without DAPI. **(E-L)** Max intensity projection of HCR detection of *Tbx6*, *Pgk*, and *Pkm* mRNA expression at 24-26ss or **(E’-L’)** midline optical sections; lateral views of PSM, **yellow arrows** highlighting posterior midline PSM expression. Control n=5, mutant n= 4. **(M)** Enzymatic lactate assay showing higher levels of lactate in the posterior (P) PSM compared to anterior (A) PSM of 24-26ss control embryos and equal amounts of lactate in mutant anterior and posterior PSM; lactate levels are normalized to control posterior, error bars represent SEM, *: p < 0.05.

To assess whether impaired vascular development contributes to the hypoxia observed in mutants, we analyzed vascularization in the PSM region at 18-20ss (Fig. S2), a stage when an increase of intracellular hypoxia has been ongoing in mutants. At this stage, no vessels are present within the PSM, but a vascular network underlies the PSM (Fig. S2), as previously reported [33]. Quantification revealed no reduction in the vascular volume near the PSM of mutants (Fig. S2I, J). By 30–32ss, vascularization within the PSM is now apparent, and mutants exhibit a significant reduction in vascular volume (Fig. S3L, M), characterized by large, poorly elaborated vessels (Fig. S3F–K). Together, these data indicate that the initial we first hypoxia observed at 10-14 ss cannot be attributed to vascular deficits, although reduced vascularization is evident at later stages, consistent with known roles of HIF1α [28].

In addition to regulating oxygen delivery, HIF-1α influences cellular oxygen consumption by upregulating glycolytic pathway genes and suppressing mitochondrial respiration [25, 32]. In mutants, the expression of key glycolytic genes, *Pgk1* and *Pkm*, is significantly reduced in the PSM and neural tube (Fig. 1I–L’, *Pgk1* reduced by 54.6% p < 0.001, *Pkm* reduced by 47.9% p < 0.001). As previously reported [19, 20], these glycolytic genes are highly expressed in the posterior PSM (Fig. 1E–H’), corresponding to the region of increased hypoxia observed in mutant embryos (Fig. 1B). This elevated posterior glycolytic gene expression generates a lactate gradient, with lactate levels higher in the posterior compared to the anterior PSM [19, 20]. While control embryos maintain this gradient, mutant embryos exhibit a flattened posterior-to-anterior lactate gradient, with posterior lactate levels reduced to those observed in the anterior PSM (Fig 1M). This suggests that cells in this region, which rely on glycolysis and lactate fermentation for energy production, require HIF1α to sustain this metabolic program. The loss of HIF1α likely forces a metabolic shift from glycolysis and lactate fermentation to mitochondrial oxidative phosphorylation, thereby increasing oxygen consumption and contributing to the observed hypoxia in these tissues.

### Mutants have defects in somitogenesis

Building on our observations of impaired glycolysis and increased intracellular hypoxia in these mutants, we explored the effects on somitogenesis and mesoderm formation. At embryonic day 18.5 (E18.5), mutants demonstrated skeletal anomalies, including abnormally shaped and anteroposteriorly (A-P) shortened vertebrae, as well as a highly penetrant extra thoracic vertebra. Out of 24 mutants examined, 21 exhibited 14-ribbed vertebrae instead of the normal 13, along with 6 lumbar vertebrae (Fig 2 A, B). The remaining 3 mutants had 14-ribbed vertebrae but only 5 lumbar vertebrae, with many of these lumbar vertebrae being hemivertebrae, suggesting that these animals also possessed an extra thoracic vertebra and a loss of lumbar vertebrae, likely not caused by a homeotic transformation. Control embryos, which experienced mild hypoxia (Fig 1A), had normal skeletons, indicating that mild hypoxia alone did not cause defects. Mutants predominantly displayed abnormally shaped vertebrae in the mid-to-lower thoracic, lumbar, and sacral regions, with the highest penetrance in the lumbar region (Fig 1C). Focusing on the lumbar region, the A-P length of lumbar vertebrae was reduced by 40% in mutants (Fig S4A, B; bracket Fig 2B). Histological sections clearly revealed the abnormal vertebral structure in mutants (Fig 2D, E), showing loss or reduction of central ossification, butterfly vertebrae (asterisks) and absence of intervertebral discs (IVD, solid yellow arrows for presence and dotted yellow arrows for absence). Cell death was observed throughout the vertebral bodies at E18.5 (Fig 2F, G), consistent with HIF1α’s known role in preventing chondrocyte death during skeletal development [34]. The observed dying cells could also be contributed from notochordal remnants that would normally give rise to a portion of the IVD, the nucleus pulposus, as HIF1α is a known survival factor for those cells as well [35].

**Fig 2.**
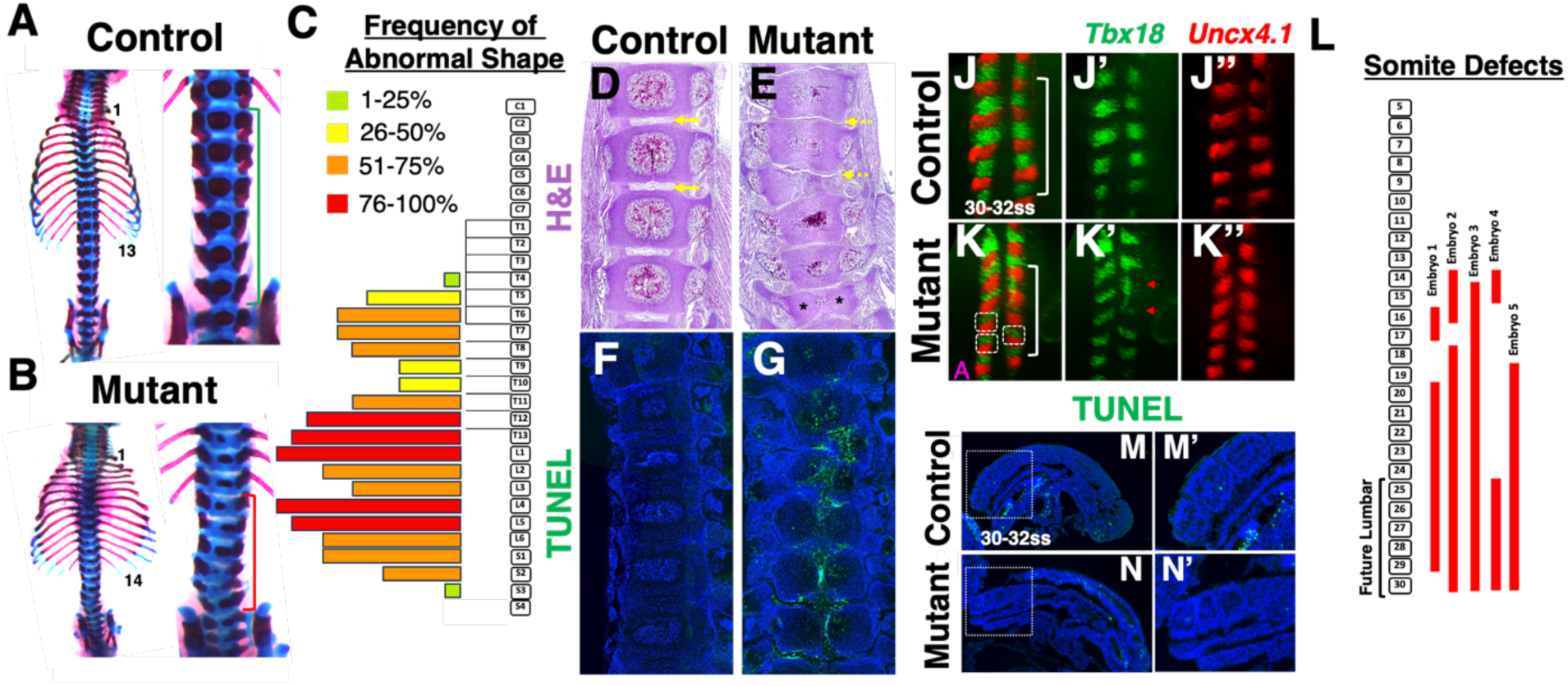
HIF1α inactivation causes vertebral and somite malformations. **(A,B)** Skeletons of E18.5 control and mutant; brackets indicate lumbar vertebrae. **(C)** Histogram showing frequency of abnormal vertebral shape along the vertebral column; n= 12, E18.5 mutant skeletons. **(D, E)** H & E staining of sections of lumbar vertebrae in E18.5 control and mutant; solid yellow arrows indicate intervertebral discs in controls and dashed arrows indicate absence of intervertebral discs in mutants, asterisks indicate hemivertebrae. **(F, G)** TUNEL staining of sections of lumbar vertebrae in E18.5 control and mutant. **(J-K’)** HCR labeling of *Tbx18* and *Uncx4.1* of somites 25-29 in 30-32ss embryos; dorsal view, red arrows indicate abnormal *Tbx18* domains, white boxes outline misaligned somites, and white brackets indicate somites 25-28. **(L)** Histogram showing range of malformations in 5 mutant embryos, each numbered box indicates somite number. **(M,N)** TUNEL staining of sections of 30-32ss control and mutant and higher magnification of somite regions **(M’,N’)** indicated by white boxes; sagittal sections, posterior to the right, dorsal up.

To assess whether vertebral abnormalities stemmed from earlier defects in somite formation, we examined somite polarity and patterning using HCR detection of *Tbx18* and *Uncx4.1*, markers for the anterior and posterior somite halves, respectively (Fig 2J, K) [31, 32]. Abnormal expression of these markers suggested defects in somite patterning, as indicated by misoriented expression stripes, particularly of *Tbx18* (red arrows, Fig 2K’), and misalignment of somites on the embryo’s left and right sides (white boxes outlining somite units, Fig 2K). Defective somites were most consistently observed in those somites giving rise to the lower thoracic and lumbar vertebrae, spanning the entire A-P range of defects observed at E18.5 (Fig 2L, bracket Fig S4D). Furthermore, somite size was significantly reduced at E9.5 (∼23% reduction, Fig S4E, F), which occurred without any increase in cell death in these somites (Fig 2M, N).

To investigate whether HIF1α was necessary within somite cells per se, we used Meox1Cre to delete HIF1α in somites but not in the PSM [36]. Meox1Cre; *Hif1α* mutants displayed a subset of the defects observed in TCre; *Hif1a* mutants, including absence of IVD, modestly A-P shortened vertebrae (∼20%), and mild alterations to the vertebral bodies (Fig S5B, E, C, F, G). However, the overall vertebral shape was not grossly affected in these mutants. Thus, while HIF1α was necessary in somites and their subsequent lineages, defective somite size and patterning likely results from segmentation defects within the PSM.

### Mesoderm formation is abnormal in mutants

Somite segmentation is governed by the interaction of two activities: the clock and the wavefront. *Tbx6* and *Msgn1* are markers of wavefront activity and are essential for keeping the PSM in an undifferentiated state [37-40]. In control embryos, *Tbx6* is expressed more broadly throughout the PSM than *Msgn1*(Fig 3A-C), extending further anteriorly and appearing earlier in cells migrating from the caudolateral epiblast (Fig 3A’’’-C’’’). Our analysis focused on the 24-26 somite stages (ss), corresponding to when somites that eventually develop into abnormal lumbar vertebrae in mutants are formed. Mutants exhibited a normal anteroposterior (AP) length and levels of *Tbx6* expression (Fig 3G, H), but had a significantly reduced volume of *Tbx6*-expressing PSM tissue (Fig 3I). This reduction was also observed at 18-20ss, when somites that will form the thoracic vertebrae are produced (Fig S6D).

**Fig 3.**
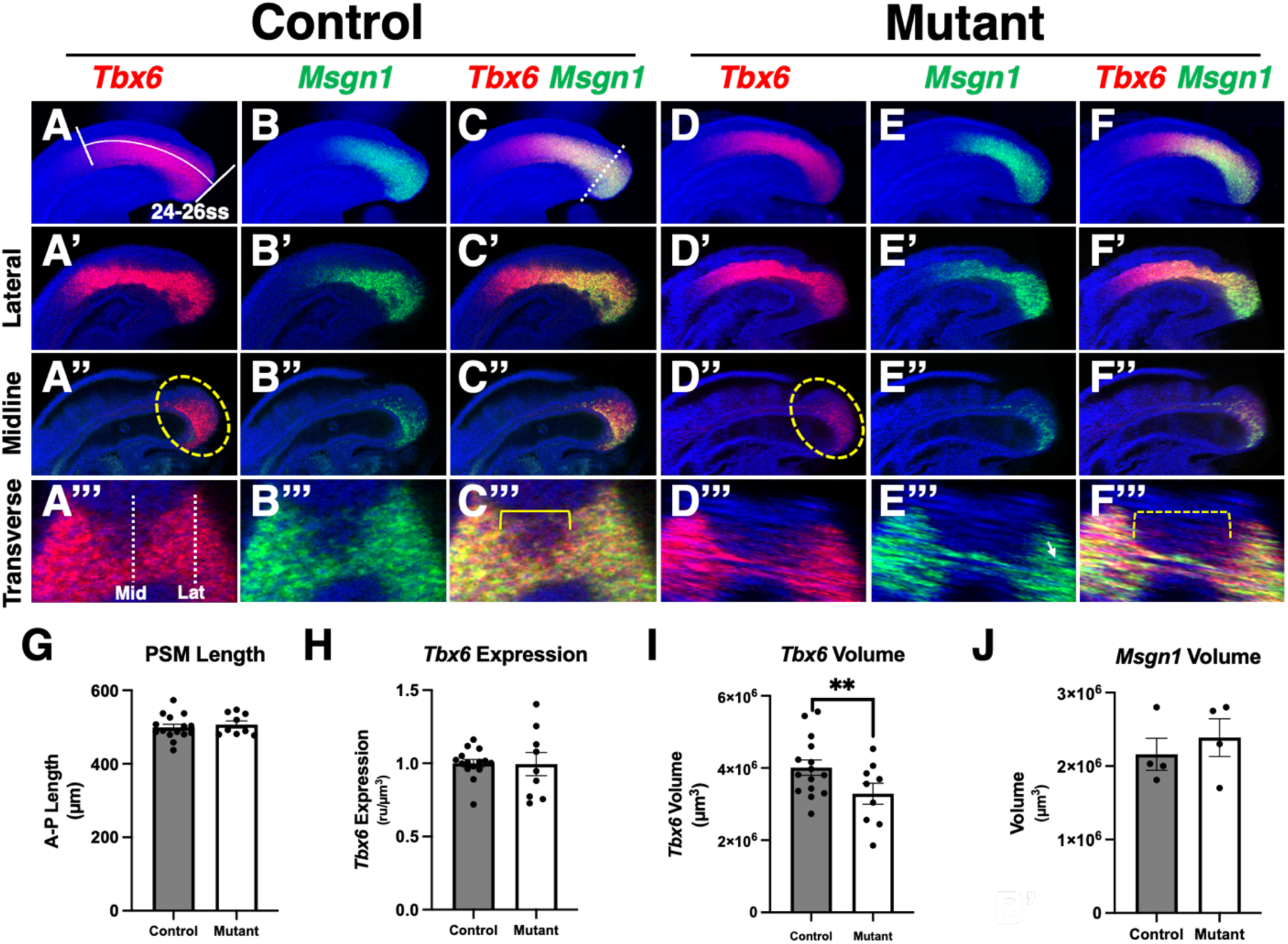
Mutants have reduced *Tbx6* volume. **(A-F)** HCR detection of *Tbx6* and *Msgn1* mRNA expression at 24-26ss in control and mutant; max intensity projection, lateral views of PSM, posterior to the right. **(A’-F’’)** Sagittal optical sections showing *Tbx6* and *Msgn1*expression at lateral (A’-F’) or midline (A’’-F’’) positions indicated by dashed lines in A’’’; lateral views of PSM, yellows circle indicates midline PSM region for comparison. **(A’’’-F’’’)** Transverse optical sections showing *Tbx6* and *Msgn1* expression in the PSM at the A-P position indicated by the dashed line in C; dotted bracket indicates region of downregulation of *Tbx6* in mutant compared to same region in control, solid bracket. **(G)** Measurement of A-P length of PSM (measurement indicated by white-bracketed line in A); no significant difference. **(H)** *Tbx6* expression within the *Tbx6* domain, normalized to *Tbx6* surface volume; no significant difference. **(I)** Volume of *Tbx6* surface; **: p < 0.01. **(J)** Volume of *Msgn1* surface; **: no significant difference.

To investigate the cause of reduced tissue volumes, we stained embryos with lysotracker-red, a marker for necrotic and apoptotic cells [36, 37]. Some cell death was noted in the PSM at later stages (32-34ss), but little was observed earlier when the reduction in *Tbx6* volume was first detected at 18-20ss (Fig S7B). Consistent with TUNEL staining (Fig 2N), abnormal levels of cell death were not found in somites using this method (Fig S7G-H). Furthermore, restoring cell survival in these mutants by removing pro-apoptotic *Bak* and *Bax* genes did not rescue the mutant phenotype (Fig S8). Although cell death was completely rescued (Fig S8C), vertebral abnormalities, including hemivertebrae, persisted (red arrows, Fig S8F, H). Consequently, cell death might be a secondary outcome of HIF1α loss, rather than a primary cause of axial skeletal defects. Instead, *Tbx6* expression analysis revealed that expression in the posterior midline PSM was absent or severely reduced (Fig 3D’’, D’’’, Fig S6B’), possibly accounting for the reduced volume. To confirm that the reduction in *Tbx6* volume did not indicate a loss of PSM, we measured the volume of *Msgn1*-expressing PSM tissue. *Msgn1* is not typically expressed in posterior midline cells, where *Tbx6* expression is lost in mutants (Fig 3B’’), and when the volume of *Msgn1*-expressing cells was measured, no difference was found between control and mutant (Fig 3J). Thus, the reduction in *Tbx6* PSM volume appears to result from a failure to initiate *Tbx6* expression in newly formed mesoderm.

The posterior midline PSM is a site where new mesoderm forms from neuromesodermal progenitors (NMPs). We assessed NMPs by examining *Sox2* and *T*, which co-express to define the NMP population (Fig 4)[41]. Mutants showed a significant decrease in *Sox2-T* double-positive tissue, indicating issues with these progenitors (Fig 4G). A similar decrease is observed when the PSM is glucose-deprived, with reduced *T* expression and increased *Sox2* expression [19]. The decline in double-positive cells in mutants is likely due to decreased *T* expression (Fig 4B”, E”, brackets). *T* is crucial for cell ingression and mesoderm program initiation [18]. A failure in *Tbx6* expression within this domain aligns with decreased *T* expression [42]. A morphological "bump" was consistently observed in mutants (arrows in Fig4E, F), suggesting issues with cells completing ingression and accumulating in the midline. While the buildup of cells might eventually resolve through cell death (Fig S7F), mutants did not lack vertebral elements (most mutants had an additional thoracic vertebra and the same number of caudal vertebrae as controls: control average was 30.4 caudal vertebrae vs 29.6 in mutants), indicating no significant mesoderm loss. Therefore, this change in ingression likely represents a delay in mesoderm identity, not a complete failure to transition to mesoderm, due to decreased *T* expression.

**Fig 4.**
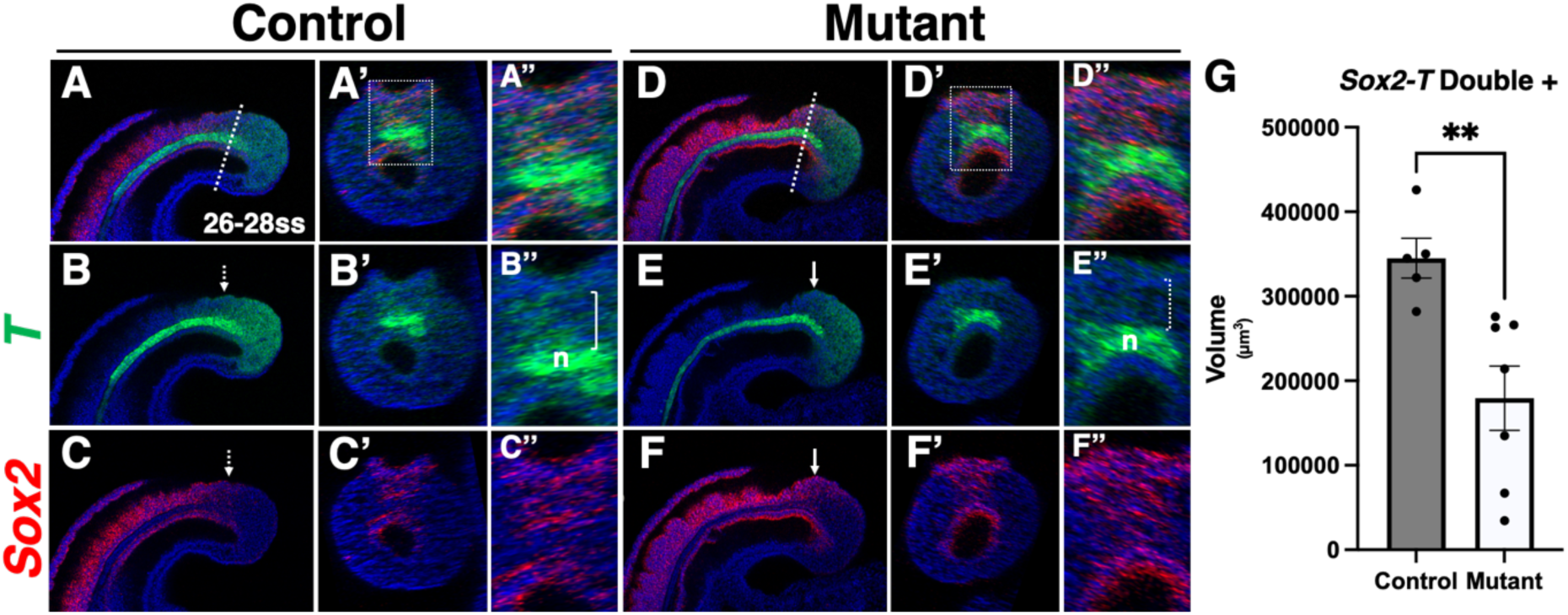
Mutants have reduced expression of *T*. **(A-F)** Midline sagittal optical sections of *T* and *Sox2* mRNA expression at 26-28ss in control and mutant embryos, and **(A’-F’)** transverse optical sections (at the position indicated by the dashed line in A or B) with **(A’’-F’’)** zoomed in regions of A’-F’ as indicated by box in A’ and D’; n: notochord, brackets in B” and E” indicate *T* expression in region of primitive streak, arrows indicate morphological bump in mutants compared to same region in controls indicated by dashed arrows. **(G)** Quantification of volume of *Sox2-T* double positive regions of the PSM in 26-28ss embryos; **: p < 0.01, error bars represent SEM.

### Wavefront is not significantly altered in mutant PSM

Wnt and FGF signaling are crucial for the maintenance of the PSM and mesoderm formation [8, 9]. *Wnt3a* expression is necessary for preserving the PSM, operating through the canonical β-catenin signaling pathway to regulate gene expression, including *Sp5*, which is positively influenced by Wnt signaling [43]. In mutants, the expression of *Wnt3a* and *Sp5*, as well as the non-canonical Wnt signaling member *Wnt5a*, remained unchanged (Fig S9)

*Fgf4* and *Fgf8* are redundantly essential for PSM maintenance, but the absence of *Fgf4* alone negatively affects segmentation clock function [5]. While *Fgf4* expression within the *Tbx6* domain was unaltered in mutants (Fig S10G), *Fgf8* expression was significantly reduced when measured within the *Tbx6* expression domain (Fig S10P). Given that the highest levels of *Fgf8* expression are located in the posterior midline of the PSM, where *Tbx6* expression is diminished in mutants, quantification within the *Tbx6* volume might not fully capture this expression domain. Therefore, we also assessed *Fgf8* expression within its own domain and confirmed a 20% reduction (Fig S10S). It is likely that this decrease in *Fgf8* expression results from previously mentioned issues with mesoderm program gene expression (Fig 4). To ascertain if the reduction of *Fgf8* affected FGF signaling in the PSM, we analyzed the expression of the FGF-responsive genes *Spry2* and *Spry4* [8]. No changes in the expression of either gene were observed in mutants (Fig S11G, H), although mutants frequently exhibited asymmetrical patterns of the anterior oscillating domain of both genes (brackets in Fig S11E’, F’; 7 out of 11 mutants showed asymmetrical expression of one or both genes). The pattern of this anterior band is influenced by Notch somite clock oscillations [44]. Thus, while FGF and Wnt signaling appear unaffected in mutant PSM, the abnormal patterns of *Spry2* and *Spry4* expression may indicate defective somite clock oscillation.

### The somite segmentation clock is abnormal in mutants

The somite segmentation clock regulates the timing and initiation of somite segmentation at the anterior boundary of the presomitic mesoderm (PSM). *Mesp2* and *Ripply2* expression trigger segmentation and establish correct somite polarity (*Tbx18* and *Uncx4.1* expression) in response to the clock [14, 45]. Normally, these genes are symmetrically expressed on the right and left sides of the embryo (Fig 5A). However, mutants often exhibited asymmetric expression patterns, with differences in expression levels and anteroposterior (A-P) position of bands (Fig 5B, 3 out of 6 mutants showed asymmetric expression). The asymmetry in *Mesp2* A-P position could explain the misalignment of somites (Fig 2K), and the asymmetry in expression levels may also cause the observed increase and decrease in *Uncx4.1* and *Tbx18*, respectively (Fig 2K, K’).

**Fig 5.**
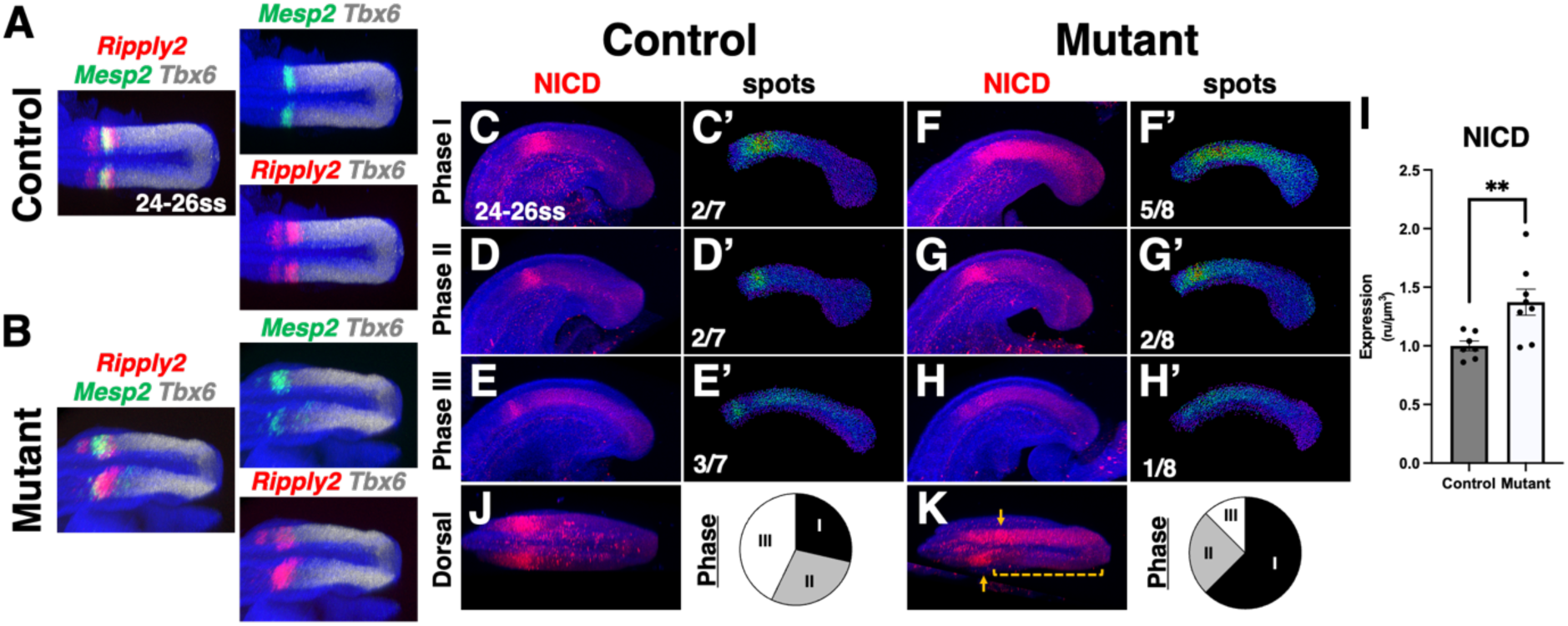
Segmentation gene and NICD oscillations are abnormal in mutants. **(A-B)** Max intensity projection of *Ripply2, Mesp2, and Tbx6* mRNA expression at 24-26ss in control and mutant; dorsal views of PSM, posterior to the right. **(C-H)** Immunostaining for NICD expression and **(C’-H’)** spot heatmaps of NICD staining at 24-26ss in control and mutant separated by oscillatory phase; numbers indicate fraction of embryos in each phase summarized in pie-charts below; lateral view, posterior to the right. **(I)** Quantification of NICD expression within the PSM; **: p < 0.01, error bars represent SEM. **(J, K)** Dorsal view of NICD showing asymmetry in mutant: bracket and arrows.

Notch signaling forms the core of the segmentation clock. Activated Notch serves as the primary clock output, patterning somite segmentation gene expression [44]. Upon binding of Notch receptors to transmembrane ligands in adjacent cells (members of the *Dll* family), the Notch intracellular domain (NICD) is cleaved and translocates to the nucleus to activate target gene expression [12]. Coordinated Notch activation among neighboring cells, followed by its repression, generates a synchronized oscillation of expression that pulses through the PSM from posterior to anterior. This oscillation is observed in fixed embryos by classifying expression patterns into oscillatory phases (I-III) and examining phase distribution [46]. In controls at 24-26ss, phase I is characterized as having a defined but wide mid- to anterior-PSM band of high NICD staining (Fig 5C, C’). In phase II, this band has moved further anterior and become more well defined (Fig 5D, D’), and in phase III this oscillation has started to dim and new less defined oscillation is building in the mid- to posterior-PSM (Fig 5E, E’). In control embryos, phase distribution was approximately one-third of embryos in each phase, consistent with normal oscillation (Fig 5C-E’) [13, 46]. In mutant embryos, NICD expression predominantly appeared as a widespread single domain in the anterior half of the PSM, resembling Phase I (Fig 5F-H’). Additionally, expression was sometimes asymmetric on the embryo’s right and left sides (Fig 5K). Lastly, NICD expression was significantly higher in mutants (Fig 5I), suggesting a possible defect in repression of Notch activation.

Segmentation clock oscillation is driven by negative feedback within the Notch pathway. *Hes7* and *Lfng* are Notch-activated genes encoding a transcriptional repressor and a glycosyltransferase, respectively, which attenuate Notch signaling [4, 44, 46]. Loss of either gene leads to desynchronization of Notch activation between cells, causing major defects in collective oscillation through the PSM. Loss of coordinated oscillation results in improperly segmented somites and, ultimately, vertebral malformations [5, 47]. The expression patterns of *Hes7* and *Lfng* can be categorized into phases akin to NICD phases (Fig S12). HES7 acts as a transcriptional repressor, inhibiting both its own expression and that of *Lfng*. As HES7 protein levels rise, *Lfng* expression is suppressed [48, 49]. This leads to an mRNA expression pattern where *Lfng* and *Hes7* expression overlap within certain regions. Over time, as HES7 protein accumulates, *Lfng* expression is abruptly repressed. *Lfng* expression peaks at the anterior end of the domain, where HES7 protein has not yet been sufficiently translated to suppress *Lfng* (brackets, Fig S12), while it diminishes posteriorly, where Hes7 has been expressed longer and has achieved higher protein levels. Notably, *Hes7* expression remains relatively elevated in the posterior even after *Lfng* repression, suggesting that *Lfng* repression requires a lower threshold of HES7 compared to the autorepression of *Hes7* expression.

At 24-26 ss, *Hes7* expression in control embryos was observed across phases I-III in nearly equal proportions (Fig 6A-C’’, F). Phase I is defined as high posterior expression, and phase II is marked by widespread *Hes7* expression throughout the presomitic mesoderm (PSM), which becomes more localized into a narrower band of expression in the anterior PSM by phase III. Mutant embryos predominantly exhibit *Hes7* expression patterns consistent with phases II and III (Fig 6D-E’’, G). In particular, in mutants *Hes7* expression remained largely throughout the PSM, although slight downregulation in the posterior was observed in some embryos (yellow arrows, Fig 6E’, E’’). This downregulation accounted for the classification of some embryos as being in phase III. However, overall, *Hes7* expression did not appear to oscillate in a coordinated manner in mutants. Moreover, *Hes7* expression levels in mutants were significantly lower than in controls.

**Fig 6.**
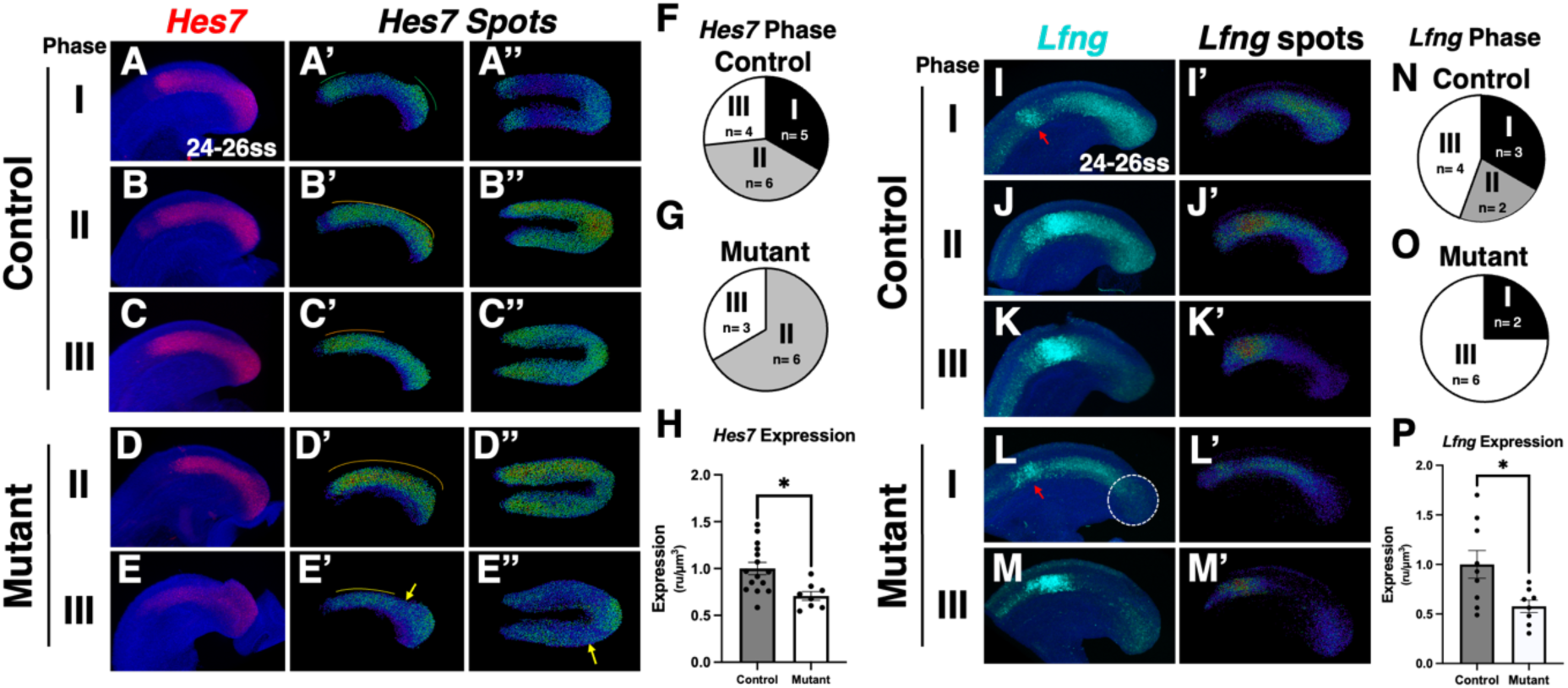
*Hes7* and *Lfng* expression and oscillations are abnormal in mutants. **(A-E)** Whole mount HCR detection and **(A’-E’’)** spot heatmaps of *Hes7* mRNA expression at 24-26ss in control and mutant; lateral views in A-E’, dorsal views in A’’-E’’, lines above images in A’-E’ indicate oscillation and are colored to reflect higher (orange) vs lower (green) intensity of band, posterior to the right. **(F, G)** Pie charts of *Hes7* phase distribution of control and mutant shown in A-E. **(H)** Quantification of *Hes7* expression within the PSM as defined by *Tbx6* expression; *: p < 0.05, error bars represent SEM. **(I-M)** Whole mount HCR detection and **(I’-M’)** spot heatmaps of *Lfng* mRNA expression at 24-26ss in control and mutant; red arrows indicate diminishing anterior band used to define phase I, and circle indicates abnormal posterior loss of expression, lateral views, posterior to the right. **(N, O)** Pie charts of *Lfng* phase distribution of control and mutant shown in I-M. **(P)** Quantification of *Lfng* expression within the PSM as defined by *Tbx6* expression; *: p < 0.05, error bars represent SEM.

Similar to *Hes7*, *Lfng* expression was aberrantly expressed, and its phase distribution was altered in mutants (Fig 6L-M’, O). Categorization of *Lfng* phases is based on similar criteria as *Hes7*, where phase I has a growing posterior domain, in addition to a thin defined band in the anterior PSM (Fig S12, Fig 6I,I’). In phase II, the posterior band of expression has propagated anteriorly and diminished posteriorly, then finally in phase III this band is thinner and at the anterior boundary of the PSM and all posterior expression is extinguished (Fig 6K, K’). Mutant embryos with a defined anterior band (red arrow, Fig 6L) and some level of posterior expression were classified as phase I; however, the observed posterior downregulation of *Lfng* in mutants (circle, Fig 6L) also suggests phase III characteristics. The majority of mutant embryos were classified as being in phase III (Fig 6M, M’, O), with overall *Lfng* expression significantly reduced (Fig 6P). This reduction is mechanistically consistent, as most mutants are classified in phase II of *Hes7* oscillation, where expanded *Hes7* expression throughout the PSM would repress *Lfng* expression in the posterior PSM. Furthermore, reduced *Lfng*, which typically modifies the Notch protein to decrease signaling, would lead to increased Notch activation as observed (Fig 5I). Therefore, the abnormal shape and patterning of vertebrae can be attributed to disrupted clock oscillations, wherein two primary negative regulators of Notch signaling, *Lfng* and *Hes7*, are misexpressed and fail to oscillate properly.

Next, we analyzed *Hes7* and *Mesp2* expression in 18-20ss embryos to investigate whether *Hes7* abnormalities manifest during the formation of the anterior somites that will develop into thoracic vertebrae, a stage when skeletal abnormalities first emerge, albeit with lower penetrance (approximately 25% of vertebrae exhibit abnormalities). At 18-20ss, the clock is much faster and therefore the characteristics of the phases are different. Phase I shows a small domain of expression beginning in the posterior PSM and a broad band in the anterior half of the PSM (Fig S13A). In phase II, the posterior band has expanded throughout the posterior half of the PSM and the anterior band is at the anterior boundary of the PSM and fading (Fig S13C). Phase III sees the loss of the anterior band and widespread expression of *Hes7* throughout most of the PSM (Fig S13E). Mutants displayed *Hes7* expression in all three phases, yet a disproportionate number were classified as phase II (Fig S13D, 7 out of 11 total embryos). Notably, a subset of mutants (4/15) exhibited asymmetric *Hes7* expression (bracket, Fig S13G). Interestingly, *Hes7* expression was not diminished at this stage (Fig S13H), suggesting that the pattern of *Hes7*, rather than its overall expression levels, contributed to segmentation defects. Furthermore, *Mesp2* expression was also asymmetrically distributed in embryos with asymmetric *Hes7* expression (Fig S13G’). We found that most mutant embryos had two bands of *Mesp2* (8/11 mutants versus 3/12 controls, Fig S13A’-F’), and most phase II mutants exhibited two *Mesp2* bands (5/7), whereas no control embryos in phase II displayed this characteristic. This finding may indicate that the oscillation clock is functioning at an accelerated rate in mutants, which aligns with the presence of extra thoracic vertebrae in these embryos.

### Increase of intracellular hypoxia is partially responsible for clock and vertebral abnormalities in mutants

To investigate whether increased hypoxia was responsible for the observed defects, we attempted to rescue PSM hypoxia by incubating pregnant dams carrying E9.0 gestating embryos in 80% oxygen (hyperoxia) for 3 hours. In all hyperoxia-treated mutants, hypoxia in the PSM and neural tube was substantially reduced compared to untreated mutants, although with some variability (Fig 7D, E). To assess the impact on skeletal patterning, we repeated the experiment, this time keeping the mice in 80% oxygen for 10 hours before returning them to normoxia and allowing development until E18.5. The gestating embryos were exposed to hyperoxia during the estimated timeframe when the somites that give rise to the lumbar vertebrae were forming (24-29ss, based on embryos generated from the same colony collected for other experiments within the incubation timeframe). Hyperoxia-treated mutants exhibited similar levels of skeletal malformations in the thoracic regions, which are formed from somites formed prior to the hyperoxia treatment window but showed significantly fewer defects in the lumbar region (Fig 7F-G). Given the variable and sometimes incomplete rescue of hypoxia in hyperoxia-treated mutants, it is reasonable to conclude that not every animal was fully rescued. Furthermore, since embryos were only exposed to hypoxia at E9.5, any later requirements in vertebral formation could not be rescued, making it unlikely the rescue of the mild vertebral defects due to this later requirement.

**Fig 7.**
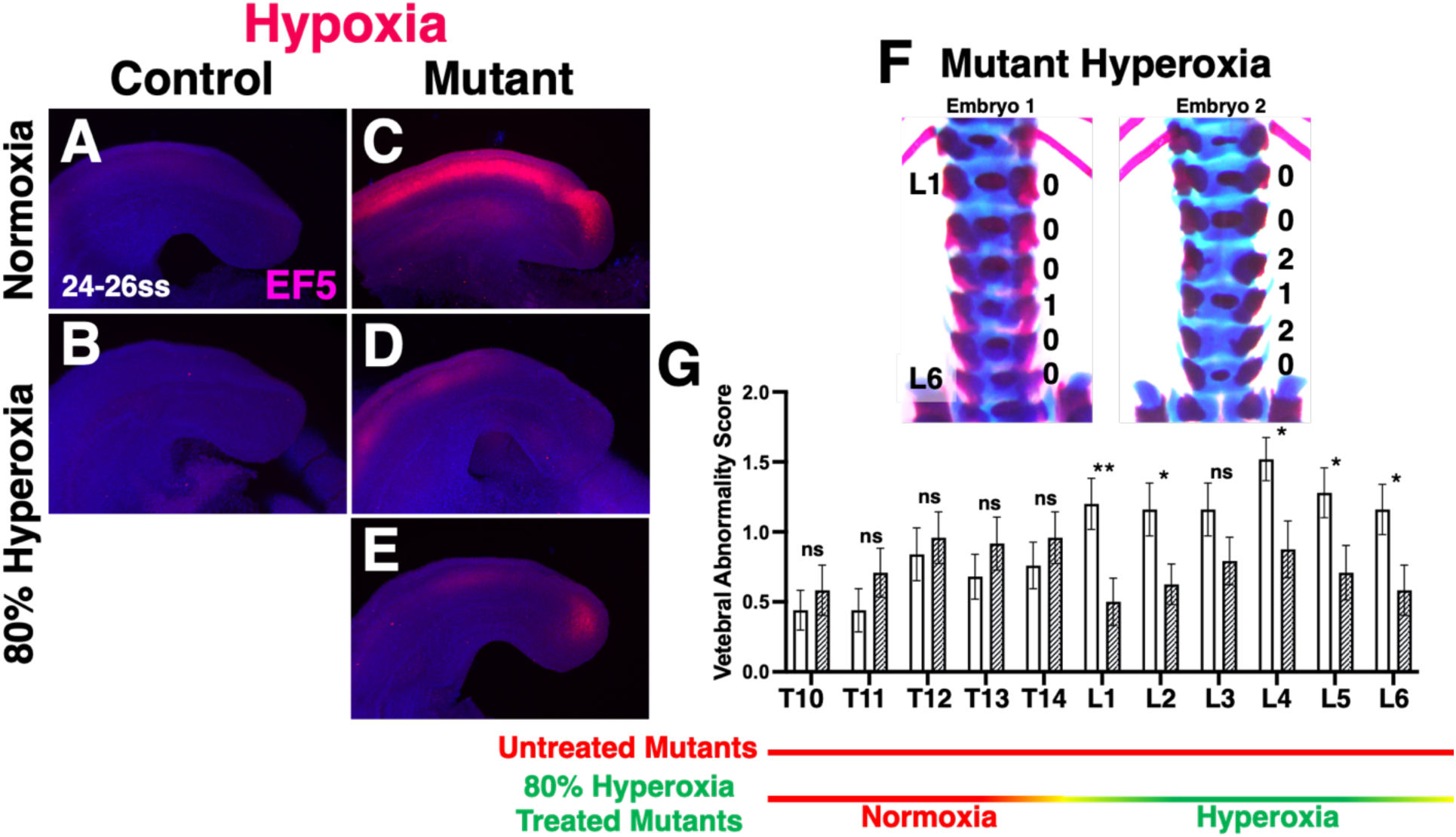
Hyperoxia partially rescues *increased intracellular* hypoxia of the PSM and vertebral malformation in mutants. **(A-E)** Whole mount EF5 detection of hypoxia in control and mutant embryos at normoxia, A and C, or exposed to gestational hyperoxia (80% Oxygen), B-E; lateral views, posterior to the right. **(F)** Skeletal preparations of hyperoxia treated mutants showing lumbar region; numbers indicate scoring of vertebral abnormalities, normal: 0, slight abnormal shape: 1, very abnormal shape: 2. **(G)** Quantification of vertebral abnormalities based on scoring system from thoracic vertebra 10 (T10) through lumbar vertebra 6 (L6); Mutant Normoxia: n = 25, Mutant Hyperoxia: n = 24, **: p < 0.05, *: p < 0.05, or ns: not significant, error bars represent SEM. Color coded lines and labels below indicate estimated timing of hyperoxia exposure.

To determine whether the rescue of skeletal formation by hyperoxia correlated with the rescue of clock and mesoderm formation defects, we repeated the hyperoxia treatment for 5 hours, followed by dissection and analysis of *Hes7* and *Tbx6* expression. Notably, *Hes7* oscillations appeared to normalize following hyperoxia treatment, with approximately one-third of embryos found in each phase, including phase I, which is never observed in mutants under normoxia (Fig 8F-H, Fig 6G). However, *Hes7* expression levels were not restored in hyperoxia-treated mutants, suggesting that the reduction in *Hes7* per se may not be responsible for the skeletal phenotype (Fig 8I). This finding aligns with our previous observation that a similar reduction of ∼25% in *Hes7*-heterozygous animals did not exceed the expression threshold necessary for normal oscillation, resulting in infrequent skeletal malformations [5].

**Fig 8.**
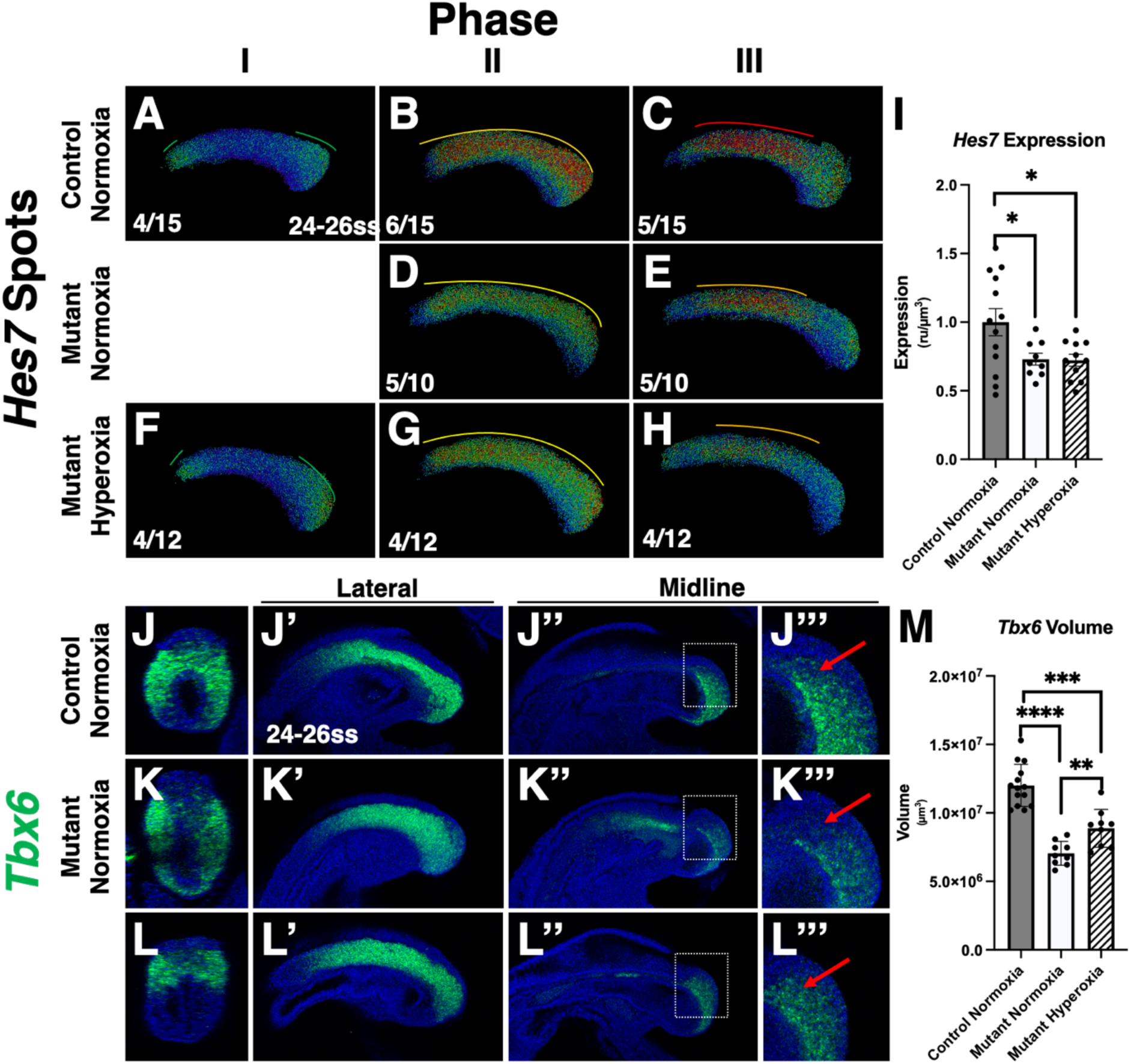
Hyperoxia partially rescues *Hes7* oscillation *Tbx6* expression in mutants. **(A-H)** Spot heatmaps of *Hes7* mRNA expression at 24-26ss in untreated (Normoxia) control and mutant or Hyperoxia treated mutant sorted by phase; numbers indicate distribution of embryos per phase, lines above images indicate oscillation and are colored to reflect higher (red) vs lower (green) intensity of band, lateral views, posterior to the right. **(I)** Quantification of *Hes7* expression within the PSM as defined by *Tbx6* expression; error bars represent SEM, *: p < 0.05 **(J-L)** Transverse or **(J’-L’’’)** sagittal optical sections of whole mount HCR detection of *Tbx6* mRNA expression at 24-26ss in control and mutant at lateral (J’-L’) or midline (J’’-L’’) planes with (J’’’-L’’’) zoomed indicated by white boxes; red arrows indicate area of rescue in hyperoxia treated mutants, lateral views of PSM, posterior to the right. **(M)** Quantification of *Tbx6* volume; ****: p < 0.0001, ***: p < 0.001,**: p < 0.01, error bars represent SEM.

*Tbx6* expression was partially rescued with hyperoxia exposure (Fig 8J-L’’’). In hyperoxia-treated mutants, posterior midline expression of *Tbx6* was increased (Fig 8L-L’’’), resulting in a significant increase in the volume of the presomitic mesoderm (PSM) compared to untreated mutants; however, this volume did not reach wild-type levels (Fig 8M). Thus, the hyperoxia-mediated rescue of defective posterior PSM formation correlates with the rescue of *Hes7* expression and the alleviation of skeletal defects. This suggests a model in which HIF1α prevents severe intracellular hypoxia in PSM progenitors that is essential for their formation, normal gene expression, and subsequent coordinated oscillation of segmentation genes.

## Discussion

Our results demonstrate that HIF1α is crucial for the normal formation of the PSM and its proper segmentation into somites, both essential steps for subsequent vertebral development. Removing Hif1α from the PSM (via TCre-mediated recombination) results in severe vertebral abnormalities, including exacerbated A-P shortening, the presence of butterfly vertebrae and hemivertebrae, which contribute to scoliosis-like curvature of the spine, as well as the appearance of an additional thoracic vertebra. In contrast, removing Hif1α later in the somitic lineage, after segmentation (with *Meox1^Cre^*), caused modest vertebral defects, that were not the result of segmentation abnormalities in the PSM and were very likely due to HIF1α’s role in chondrocyte survival per se [34]. Notably, both Meox1^Cre^- and TCre-mediated deletion of *Hif1α* resulted in the loss of intervertebral discs, due to a requirement for HIF1α in this tissue [50]. Thus we conclude that HIF1α has a unique and non-redundant role in the PSM.

In the PSM of mutant embryos, there is an increase in intracellular hypoxia and downregulation of glycolytic enzymes. Notably, at earlier stages, when hypoxia is increased, the vascular network surrounding the PSM is normal, and becomes abnormal only at later developmental stages. Since intracellular hypoxia is a net result of oxygen availability and consumption, our findings suggest that the increase in intracellular hypoxia during the earlier stages of PSM development is due to increased oxygen consumption that occurs in the absence of HIF1α, which leads to heightened mitochondrial respiration. These findings are consistent with the observation that the PSM is highly glycolytic and that HIF1α plays a critical role in reprogramming metabolism by upregulating glycolysis while suppressing mitochondrial oxygen consumption [20, 21, 24, 30, 32, 51]. This is consistent with our recent report that in the developing chondrocytic growth plate, which gives rise to most of the vertebrate skeleton, the loss of HIF1α increases intracellular hypoxia by upregulating mitochondrial respiration. This increase in intracellular hypoxia leads to delayed chondrocyte differentiation from mesenchymal precursors [32]. This suggests the possibility that heightened intracellular hypoxia in our mutants causes the delay of mesoderm gene expression including *Tbx6*. This idea is supported by the rescue of both the defective posterior PSM formation and *Tbx6* expression in mutants exposed to conditions that restores oxygenation of the PSM.

Abnormal clock oscillations account for most vertebral malformations observed in mutants. The asymmetric vertebral shapes and presence of hemivertebrae in these mutants can be attributed to asymmetric *Mesp2* patterns generated by asymmetric NICD activation and *Hes7* expression. Furthermore, the timing and frequency of *Hes7* oscillation loss correlate with the incidence of malformations. In 18-20ss mutant embryos, all three phases of *Hes7* oscillatory behavior are present, although phase II is disproportionately represented. The somites and subsequent vertebrae generated by these oscillations exhibit abnormalities approximately 25% of the time. By 24-26ss, the mutant clock oscillations are consistently abnormal, as are the associated somite and vertebral defects. *Hes7* represses both itself and *Lfng*, while *Lfng* attenuates Notch activation. The failure of normal *Hes7* oscillation, resulting in widespread *Hes7* expression, explains the diminished and overall lack of posterior *Lfng* expression. The downregulation of *Lfng* leads to an increase in NICD levels. Although widespread expression of *Hes7* should induce its autorepression, the loss of HIF1α appears to disrupt this autorepression loop, likely leading to the segmentation clock abnormalities observed in mutants.

This is the first loss-of-function genetic study charactering a phenotype in which both glycolysis and Notch clock oscillations are disrupted in the presomitic mesoderm. As such, our *in vivo* data is supported by and complements several recent studies, based on cultured pluripotent stem cell-derived models or embryonic explants, that analyze the dynamic relationship between metabolism and the somitic clock [19, 20, 52-55]. Several of these studies demonstrate that the period of clock oscillations scale with glycolytic flux [54, 55]. Incubation with non-glycolytic substrates in these *in vitro* models causes effects that phenocopy aspects of our mutant defects: loss of *Lfng* expression in the posterior PSM [20], or a reduced expression of *Hes7* [54]. However, the role of glycolysis may extend beyond a bio-energetic function to provide signaling metabolites that regulate oscillations [52, 53]. Thus, the reduced glycolytic activity we observe may contribute to the mutant phenotype; however, our observation that both mutant clock oscillations and vertebral malformations are substantially rescued during gestational hyperoxia indicates that oxygen levels *per se* may a central cause of these defects.

The next critical question is how the increase in intracellular hypoxia due to the loss of HIF1α alters clock oscillations and somite segmentation. Loss of HIF1α in the presomitic mesoderm (PSM) increases cell death at this site, which is consistent with the augmented cell death observed in other developmental contexts, such as the developing growth plate and the nucleus pulposus. However, this increase in cell death is likely not the primary cause of the observed effects, as it occurs only at late stages of development when the abnormal PSM/somite phenotype is already established.

*Hes7* and *Lfng* have been shown to be negatively affected by gestational hypoxia, potentially through alterations in FGF signaling [2]. However, we do not observe any changes in FGF-reporter expression or in *Fgf4*, which is critical for clock oscillation [5].

Notably, in addition to mitochondrial respiration and control of HIF stability and transcriptional activity, O2 is an essential substrate for a variety of enzymatic reactions, many of which modify the epigenetic landscape and, thus, the cellular transcriptome [56, 57]. For example, the Ten-eleven translocation proteins, which catalyze oxidation of 5-methylcytosine, and the Jumonji C domain histone lysine de-methylases use oxygen as a substrate [56]. More studies are warranted to understand how the increase in intracellular hypoxia following the loss of HIF1α affects somitogenesis.

The later onset of segmentation defects in mutants may be attributed to several factors. The defects observed in mutants coincide with the approximate timing of placental labyrinth formation and maternal gas exchange [58]. At this stage of development, rodents undergo a significant shift from glycolysis and lactate fermentation to oxidative phosphorylation [59]. HIF1α may be essential for suppressing oxidative phosphorylation during this transition to protect the PSM from severe hypoxia. Additionally, there is a gradual slowing of somite formation around 18 ss. On day E8.5, somites are formed at a rate of nearly one per hour; however, over the following 24 hours, this rate slows to approximately one every two hours [60]. We observe evidence of this in 18ss control embryos, which exhibit 1-2 bands of *Mesp2* expression, while 24-26ss controls consistently show only a single band of *Mesp2*, indicative of a slower rate of somite formation. Conversely, we find evidence of a faster rate of somite formation in mutants at 18ss, as 8 out of 11 mutants displayed two *Mesp2* bands, and adults had an extra thoracic vertebra. It is possible that the loss of HIF1α leads to an increase in clock oscillation rate, which may have minimal effects during early somitogenesis when clock speed is typically faster. However, this increase could detrimentally impact somite formation when the clock speed decreases to accommodate the formation of larger lumbar vertebrae. Recent studies have shown that glycolysis affects segmentation clock speed, with lower glycolytic flux corresponding to a faster clock speed [61]. Therefore, it is possible that the reprogramming of metabolism, i.e. reduction of glycolysis and increase of oxidative phosphorylation upon loss of HIF1α, may contribute to speed up somitogenesis. Finally, the formation of smaller somites may also be explained by a scaling mechanism, where a reduction in *Tbx6* volume translates into the production of smaller somites [62].

In conclusion, our findings highlight the critical role of HIF1α in the formation of the PSM and its segmentation into somites, which are essential for proper vertebral development. The loss of HIF1α leads to significant vertebral abnormalities characterized by shortened and misshapen vertebrae, particularly when HIF1α is deleted from the PSM. This study reveals that increased intracellular hypoxia and altered metabolic pathways, particularly the shift from glycolysis to oxidative phosphorylation, significantly impact somite formation and the segmentation clock. Moreover, abnormal clock oscillations, evidenced by disrupted *Hes7* and *Lfng* expression, correlate with the observed malformations in HIF1α mutants. Understanding these mechanisms provides insight into the complex interplay between metabolism, gene expression, and developmental processes, suggesting potential therapeutic avenues for addressing congenital vertebral malformations.

## Methods and Materials

### Animal Use Statement

All research utilizing live mice was carried out in accordance with the recommendations in the Guide for the Care and Use of Laboratory Animals (National Academies Press; 8^th^ edition), and all experiments were covered by a protocol approved by the NCI-Frederick IACUC.

### Mouse Lines and Breeding

Mice were kept on a mixed background, with the most wildtype littermates serving as controls. Crosses were performed wherein the male always carried the Cre allele and females the *Hif1α-Flox* allele. The following alleles were used and genotyped as previously described: TCre [31], *Hif1α* -*flox* and *Hif1α -null* [34], Meox1Cre [36], *Bak* and *Bax* [6, 63]. Wild-type NIH Swiss mice were purchased from Charles River (code 550).

### HCR and Clearing

HCR was performed as previously described [5]. V3.0 split-initiator probes were purchased from Molecular Instruments. Embryos were incubated overnight in 0.5 µg/mL DAPI in PBTx (1% Triton-X100 in PBS), then mounted in agarose and cleared as previously described [5].

### Imaging

Cleared embryos were imaged on a Nikon A1R confocal, using UPlanApo 10x objective (NA: 0.40), with an image size of 1024x1024 pixels, using 405nm, 488nm, 561nm, 647nm, and 730nm lasers, to excite DAPI, Alexa Fluor 488, Alexa Fluor 546, Alexa Fluor 647, and Alexa Fluor 750, respectively. All images are Max Intensity Projections.

### H&E and TUNEL staining

For routine histology and H&E staining, tissue was fixed in 4% PFA/PBS and processed as previously reported [64]. TUNEL assay was performed using an In Situ Cell Death Detection kit (Roche) according to the manufacturer’s conditions on paraffin sections [64].

### Skeletal Preparations, Lysotracker Staining, and NICD Staining

Skeletal preparations were performed as previously described (ref-manipulating the mouse embryo). Lysotracker staining was performed as previously described [65] with the change of using 1 µL per mL of Lysotracker solution. Activated Notch (NICD) staining was performed as previously described [5].

### Measurements, Graphs, and Statistics

Vertebral length measurements were made on images that were taken at the same magnification and with a ruler on the stage, then measured in Fiji [66] and converted to mm. All graphs were made in Prism 10 (GraphPad) and a Students two-tailed T-test was used to calculate pairwise significance.

### Recombination Efficiency QPCR

Sectioned PSM was scraped into tubes using clean pipette tips. 5 wild type control samples and 4 TCre; *Hif1α ^flox/null^* samples were used for this analysis. Genomic DNA was then isolated using a QIAamp DNA FFPE Tissue Kit (Qiagen # 56404). DNA was dissolved in 20µL of ATE buffer then QPCR was done using Sybr Green and the following primers to detect *Hif1α*: 5’-TGA TGT GGG TGC TGG TGT C-3’, 5’-TTG TGT TGG GGC AGT ACT G-3’; and *Beta2-Mircoglobulin*: 5′-TCATTAGGGAGGAGCCAATG-3′, 5′-ATCCCCTTTCGTTTTTGCTT-3′. The delta-delta CT method was used to calculate relative recombination [67].

### EF5 Hypoxia Detection

Pregnant dams were injected with 10µL/g-body weight EF5 solution (Sigma SML1961-25MG, 10mM diluted in 0.9% sterile injectable saline) and then sacrificed 3 hours later. Embryos were dissected in cold PBS, fixed for 2 hours at 4°C, then stepwise dehydrated into 100% methanol and stored at -20°C until ready to stain. Embryos were stepwise rehydrated into PBT (PBS + 0.1% tween) then blocked in PBTx (PBS + 1% Triton X-100) with 10% FCS for 2 hours. Anti-EF5-Cy5 direct conjugate antibody (EMD Millipore Corp, EF5011-250UG) was diluted 1:500 in block and rocked for 4 hours at RT, then overnight at 4°C. Embryos were then washed in PBT 3 x 20’, counterstained with DAPI (0.5 µg/mL DAPI in PBTx) overnight rocking at RT, then embedded and cleared as previously described [5].

### Hyperoxia

Hyperoxia experiments were done using a Biospherix Oxycycler A41. Pregnant dams carrying E9.5 *Hif1α* mutant and control embryos were ramped up to 80% oxygen over the course of 30 minutes, then kept at 80% for 3, 5, or 10 hours before being immediately euthanized and placed in fixation for the shorter incubations or returned to cages and allowed to develop to E18.5 for the 10 hour incubation. For EF5 staining, pregnant dams were first incubated at 80% oxygen for an hour, then removed and injected with EF5 as described above and quickly returned to the chamber and incubated for two more hours, for a total of 3 hours at 80% oxygen.

### Imaris Modeling

Imaris software (Bitplane Inc, V10.0.1) was used extensively to model and quantify a variety of different features throughout this paper as follows.

<u>Surfaces</u> were used throughout to quantify expression within a volume of tissue defined by expression of a given tissue marker. The markers used to generate surfaces are indicated in each figure legend. Surface creation parameters were as previously described [5].

<u>Spot models</u> of expression were made as previously described [5], with the change that no PSF modeling was done and thresholds varied between experiments. Thresholds were chosen for a given experiment based on expression levels within controls, then held constant for all modeling in controls and mutants.

*Pecam* volume measurements were done using a combination of spot and surface models. For 18-20ss (Fig S2), a *Tbx6* surface (Fig S2 B) and *Pecam* spot model (Fig S2 C) was made first. Then a Matlab plugin was used to define *Pecam* spots that were within 75µm of the *Tbx6* surface (Fig S2 D, E). The *Pecam* signal was then masked within the *Tbx6*-near spots (Fig S2 E) and this masked signal was used to create a surface (Fig S2 F) and the volume of this surface was measured. For *Pecam* modeling at 30-32ss (Fig S3), a Tbx6 surface was made (Fig S3 C), then *Pecam* expression was masked within the surface (Fig S3 D), then a surface was made from that masked expression (Fig S3 E) and the volume of this surface was measured.

Somite A-P length measurements were taken from 30-32ss embryos in which *Uncx4.1* and *Tbx18* had been labeled by HCR and DAPI counterstained, then confocal imaged. A surface was made from the DAPI channel, and “Measurement Points” bound to the DAPI surface was used to measure the distance between anterior sides of *Uncx4.1* expression.

### Lactate Assay

Embryos were dissected in cold PBS and the PSM from each was isolated by cutting at the boundary between the first somite and the PSM. The PSM fragment was then cut in half, separating the anterior and posterior halves. These fragments were snap frozen in a slurry of crushed dry ice and ethanol then stored at -80°C. Lactate levels were measured using the Lactate Assay Kit from BioVision (K607-100) as per kit instructions using 50µL total volume for the assay and measuring OD at 570nm.

## Acknowledgments

We would like to thank L. Ostendorf-Snell, Q. Fan, for technical assistance with this project.

## Funding

This work was supported by the Intramural Research Program of the National Institutes of Health, National Cancer Institute, Center for Cancer Research (M.A., M. L) and by NIH R01 HD112003 to E.S.

## Author Contributions

*Conceptualization*: M.A., A.Y., S.S., M.L. *Writing of manuscript*: M.A., S.S., M.L. *Performed Experiments*: M.A., A.Y., B.L. All authors have reviewed and approve of the submitted manuscript.

## Competing Interest

The authors declare that they have no competing interests." If this is not accurate, please list the competing interests. All data needed to evaluate the conclusions in the paper are present in the paper and/or the Supplementary Materials

## Notes

### Competing Interest Statement

The authors have declared no competing interest.

## References

1. Acloque, H., et al., Reciprocal repression between Sox3 and snail transcription factors defines embryonic territories at gastrulation. Dev Cell, 2011. 21(3): p. 546–58.

2. Sparrow, D.B., et al., A mechanism for gene-environment interaction in the etiology of congenital scoliosis. Cell, 2012. 149(2): p. 295–306.

3. Kotch, L.E., et al., Defective vascularization of HIF-1alpha-null embryos is not associated with VEGF deficiency but with mesenchymal cell death. Dev Biol, 1999. 209(2): p. 254–67.

4. Evrard, Y.A., et al., lunatic fringe is an essential mediator of somite segmentation and patterning. Nature, 1998. 394(6691): p. 377-81.

5. Anderson, M.J., et al., Fgf4 maintains Hes7 levels critical for normal somite segmentation clock function. Elife, 2020. 9.

6. Anderson, M.J., T. Schimmang, and M. Lewandoski, An FGF3-BMP Signaling Axis Regulates Caudal Neural Tube Closure, Neural Crest Specification and Anterior-Posterior Axis Extension. PLoS Genet, 2016. 12(5): p. e1006018.

7. Young, T., et al., Cdx and Hox genes differentially regulate posterior axial growth in mammalian embryos. Dev Cell, 2009. 17(4): p. 516–26.

8. Naiche, L.A., N. Holder, and M. Lewandoski, FGF4 and FGF8 comprise the wavefront activity that controls somitogenesis. Proc Natl Acad Sci U S A, 2011. 108(10): p. 4018–23.

9. Dunty, W.C., Jr., et al., Wnt3a/beta-catenin signaling controls posterior body development by coordinating mesoderm formation and segmentation. Development, 2008. 135(1): p. 85–94.

10. Wahl, M.B., et al., FGF signaling acts upstream of the NOTCH and WNT signaling pathways to control segmentation clock oscillations in mouse somitogenesis. Development, 2007. 134(22): p. 4033–41.

11. Cooke, J. and E.C. Zeeman, A clock and wavefront model for control of the number of repeated structures during animal morphogenesis. J Theor Biol, 1976. 58(2): p. 455–76.

12. Lewis, J., A. Hanisch, and M. Holder, Notch signaling, the segmentation clock, and the patterning of vertebrate somites. J Biol, 2009. 8(4): p. 44.

13. Bone, R.A., et al., Spatiotemporal oscillations of Notch1, Dll1 and NICD are coordinated across the mouse PSM. Development, 2014. 141(24): p. 4806-16.

14. Saga, Y., et al., Mesp2: a novel mouse gene expressed in the presegmented mesoderm and essential for segmentation initiation. Genes Dev, 1997. 11(14): p. 1827–39.

15. Bardot, E.S. and A.K. Hadjantonakis, Mouse gastrulation: Coordination of tissue patterning, specification and diversification of cell fate. Mech Dev, 2020. 163: p. 103617.

16. Cano, A., et al., The transcription factor snail controls epithelial-mesenchymal transitions by repressing E-cadherin expression. Nat Cell Biol, 2000. 2(2): p. 76–83.

17. Arnold, S.J., et al., Pivotal roles for eomesodermin during axis formation, epithelium-to-mesenchyme transition and endoderm specification in the mouse. Development, 2008. 135(3): p. 501–11.

18. Wilson, V., et al., The T gene is necessary for normal mesodermal morphogenetic cell movements during gastrulation. Development, 1995. 121(3): p. 877–86.

19. Oginuma, M., et al., A Gradient of Glycolytic Activity Coordinates FGF and Wnt Signaling during Elongation of the Body Axis in Amniote Embryos. Dev Cell, 2017. 40(4): p. 342–353 e10.

20. Bulusu, V., et al., Spatiotemporal Analysis of a Glycolytic Activity Gradient Linked to Mouse Embryo Mesoderm Development. Dev Cell, 2017. 40(4): p. 331–341 e4.

21. Diaz-Cuadros, M., et al., Author Correction: Metabolic regulation of species-specific developmental rates. Nature, 2023. 616(7956): p. E4.

22. Riester, M., et al., The Warburg effect: persistence of stem-cell metabolism in cancers as a failure of differentiation. Ann Oncol, 2018. 29(1): p. 264–270.

23. Vander Heiden, M.G., L.C. Cantley, and C.B. Thompson, Understanding the Warburg effect: the metabolic requirements of cell proliferation. Science, 2009. 324(5930): p. 1029-33.

24. Kierans, S.J. and C.T. Taylor, Regulation of glycolysis by the hypoxia-inducible factor (HIF): implications for cellular physiology. J Physiol, 2021. 599(1): p. 23–37.

25. Semenza, G.L., et al., Transcriptional regulation of genes encoding glycolytic enzymes by hypoxia-inducible factor 1. J Biol Chem, 1994. 269(38): p. 23757–63.

26. Li, H.S., et al., HIF-1alpha protects against oxidative stress by directly targeting mitochondria. Redox Biol, 2019. 25: p. 101109.

27. Taylor, C.T. and C.C. Scholz, The effect of HIF on metabolism and immunity. Nat Rev Nephrol, 2022. 18(9): p. 573–587.

28. Sparrow, D.B., G. Chapman, and S.L. Dunwoodie, The mouse notches up another success: understanding the causes of human vertebral malformation. Mamm Genome, 2011. 22(7-8): p. 362–76.

29. Ducsay, C.A., et al., Gestational Hypoxia and Developmental Plasticity. Physiol Rev, 2018. 98(3): p. 1241–1334.

30. Iyer, N.V., et al., Cellular and developmental control of O2 homeostasis by hypoxia-inducible factor 1 alpha. Genes Dev, 1998. 12(2): p. 149–62.

31. Perantoni, A.O., et al., Inactivation of FGF8 in early mesoderm reveals an essential role in kidney development. Development, 2005. 132(17): p. 3859–71.

32. Yao, Q., et al., Suppressing Mitochondrial Respiration Is Critical for Hypoxia Tolerance in the Fetal Growth Plate. Dev Cell, 2019. 49(5): p. 748–763 e7.

33. Walls, J.R., et al., Three-dimensional analysis of vascular development in the mouse embryo. PLoS One, 2008. 3(8): p. e2853.

34. Schipani, E., et al., Hypoxia in cartilage: HIF-1alpha is essential for chondrocyte growth arrest and survival. Genes Dev, 2001. 15(21): p. 2865–76.

35. Merceron, C., et al., Loss of HIF-1alpha in the notochord results in cell death and complete disappearance of the nucleus pulposus. PLoS One, 2014. 9(10): p. e110768.

36. Jukkola, T., et al., Meox1Cre: a mouse line expressing Cre recombinase in somitic mesoderm. Genesis, 2005. 43(3): p. 148–53.

37. Chapman, D.L. and V.E. Papaioannou, Three neural tubes in mouse embryos with mutations in the T-box gene Tbx6. Nature, 1998. 391(6668): p. 695-7.

38. Nowotschin, S., et al., Interaction of Wnt3a, Msgn1 and Tbx6 in neural versus paraxial mesoderm lineage commitment and paraxial mesoderm differentiation in the mouse embryo. Dev Biol, 2012. 367(1): p. 1-14.

39. Chalamalasetty, R.B., et al., The Wnt3a/beta-catenin target gene Mesogenin1 controls the segmentation clock by activating a Notch signalling program. Nat Commun, 2011. 2: p. 390.

40. Oginuma, M., et al., Mesp2 and Tbx6 cooperatively create periodic patterns coupled with the clock machinery during mouse somitogenesis. Development, 2008. 135(15): p. 2555–62.

41. Koch, F., et al., Antagonistic Activities of Sox2 and Brachyury Control the Fate Choice of Neuro-Mesodermal Progenitors. Dev Cell, 2017. 42(5): p. 514–526 e7.

42. Hofmann, M., et al., WNT signaling, in synergy with T/TBX6, controls Notch signaling by regulating Dll1 expression in the presomitic mesoderm of mouse embryos. Genes Dev, 2004. 18(22): p. 2712–7.

43. Fujimura, N., et al., Wnt-mediated down-regulation of Sp1 target genes by a transcriptional repressor Sp5. J Biol Chem, 2007. 282(2): p. 1225–37.

44. Ferjentsik, Z., et al., Notch is a critical component of the mouse somitogenesis oscillator and is essential for the formation of the somites. PLoS Genet, 2009. 5(9): p. e1000662.

45. Takahashi, Y., et al., Mesp2 initiates somite segmentation through the Notch signalling pathway. Nat Genet, 2000. 25(4): p. 390–6.

46. Bessho, Y., et al., Dynamic expression and essential functions of Hes7 in somite segmentation. Genes Dev, 2001. 15(20): p. 2642–7.

47. Okubo, Y., et al., Lfng regulates the synchronized oscillation of the mouse segmentation clock via trans-repression of Notch signalling. Nat Commun, 2012. 3: p. 1141.

48. Bessho, Y., et al., Periodic repression by the bHLH factor Hes7 is an essential mechanism for the somite segmentation clock. Genes Dev, 2003. 17(12): p. 1451–6.

49. Chen, J., L. Kang, and N. Zhang, Negative feedback loop formed by Lunatic fringe and Hes7 controls their oscillatory expression during somitogenesis. Genesis, 2005. 43(4): p. 196–204.

50. Silagi, E.S., et al., The role of HIF proteins in maintaining the metabolic health of the intervertebral disc. Nat Rev Rheumatol, 2021. 17(7): p. 426–439.

51. Carmeliet, P., et al., Role of HIF-1alpha in hypoxia-mediated apoptosis, cell proliferation and tumour angiogenesis. Nature, 1998. 394(6692): p. 485-90.

52. Miyazawa, H., et al., A noncanonical role of glycolytic metabolites controlling the timing of mouse embryo segmentation. Sci Adv, 2025. 11(38): p. eadz9606.

53. Miyazawa, H., et al., Glycolytic flux-signaling controls mouse embryo mesoderm development. Elife, 2022. 11.

54. Diaz-Cuadros, M., et al., Metabolic regulation of species-specific developmental rates. Nature, 2023. 613(7944): p. 550-557.

55. Matsuda, M., J. Lazaro, and M. Ebisuya, Metabolic activities are selective modulators for individual segmentation clock processes. Nat Commun, 2025. 16(1): p. 845.

56. McDonough, M.A., et al., Structural studies on human 2-oxoglutarate dependent oxygenases. Curr Opin Struct Biol, 2010. 20(6): p. 659–72.

57. Chakraborty, A.A., et al., Histone demethylase KDM6A directly senses oxygen to control chromatin and cell fate. Science, 2019. 363(6432): p. 1217-1222.

58. Woods, L., V. Perez-Garcia, and M. Hemberger, Regulation of Placental Development and Its Impact on Fetal Growth-New Insights From Mouse Models. Front Endocrinol (Lausanne), 2018. 9: p. 570.

59. Shepard, T.H., T. Tanimura, and H.W. Park, Glucose absorption and utilization by rat embryos. Int J Dev Biol, 1997. 41(2): p. 307–14.

60. Tam, P.P., The control of somitogenesis in mouse embryos. J Embryol Exp Morphol, 1981. 65 Suppl: p. 103-28.

61. Miyazawa, H., et al., Glycolysis–Wnt signaling axis tunes developmental timing of embryo segmentation. bioRxiv, 2024: p. 2024.01.22.576629.

62. Lauschke, V.M., et al., Scaling of embryonic patterning based on phase-gradient encoding. Nature, 2013. 493(7430): p. 101-5.

63. Takeuchi, O., et al., Essential role of BAX,BAK in B cell homeostasis and prevention of autoimmune disease. Proc Natl Acad Sci U S A, 2005. 102(32): p. 11272–7.

64. Mangiavini, L., C. Merceron, and E. Schipani, Analysis of Mouse Growth Plate Development. Curr Protoc Mouse Biol, 2016. 6(1): p. 67–130.

65. Fogel, J.L., T.Z. Thein, and F.V. Mariani, Use of LysoTracker to detect programmed cell death in embryos and differentiating embryonic stem cells. J Vis Exp, 2012(68).

66. Schindelin, J., et al., Fiji: an open-source platform for biological-image analysis. Nat Methods, 2012. 9(7): p. 676-82.

67. Livak, K.J. and T.D. Schmittgen, Analysis of relative gene expression data using real-time quantitative PCR and the 2(-Delta Delta C(T)) Method. Methods, 2001. 25(4): p. 402–8.

