## Supplemental Figures for "HIF1α controls somitogenesis and spine development by regulating levels of intracellular oxygen in the presomitic mesoderm": Supplemental Figures.pdf

### Supplemental Figure Legends

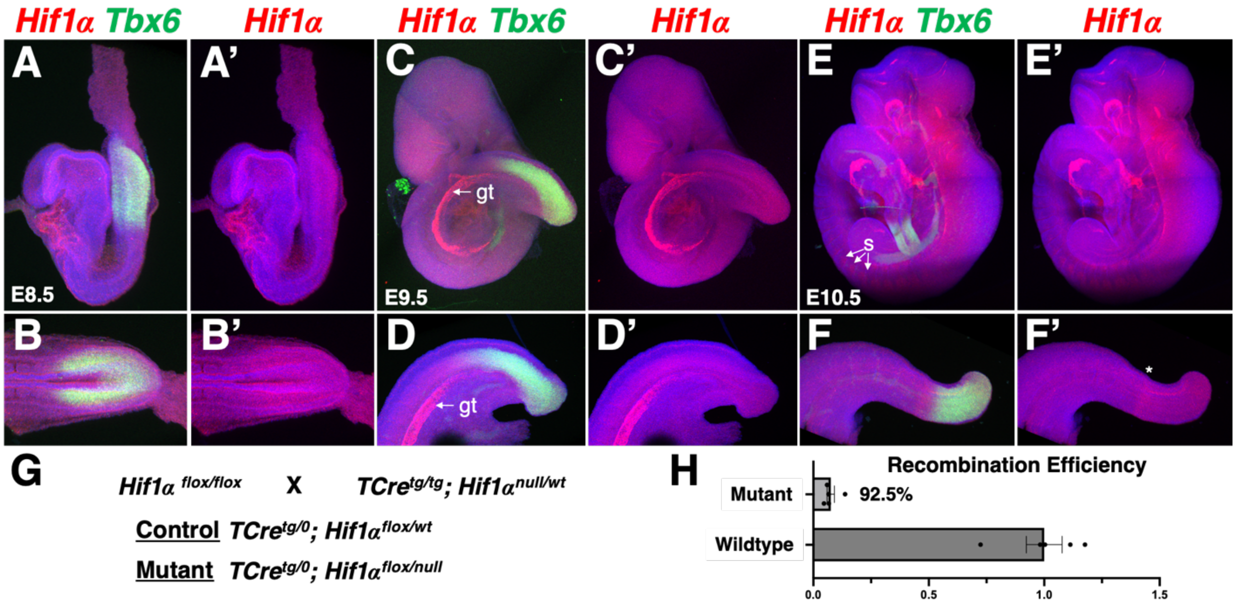


**Fig S1. *Hif1α* expression and conditional inactivation. (A-F')** HCR detection of *Hif1a* and *Tbx6* mRNA expression at stages indicated; lateral views A-E', D-F', dorsal view B, B', gt= gut tube, s = somites. **(G)** *Hif1a* inactivation breeding scheme. **(H)** Quantification of *Hif1a* recombination.

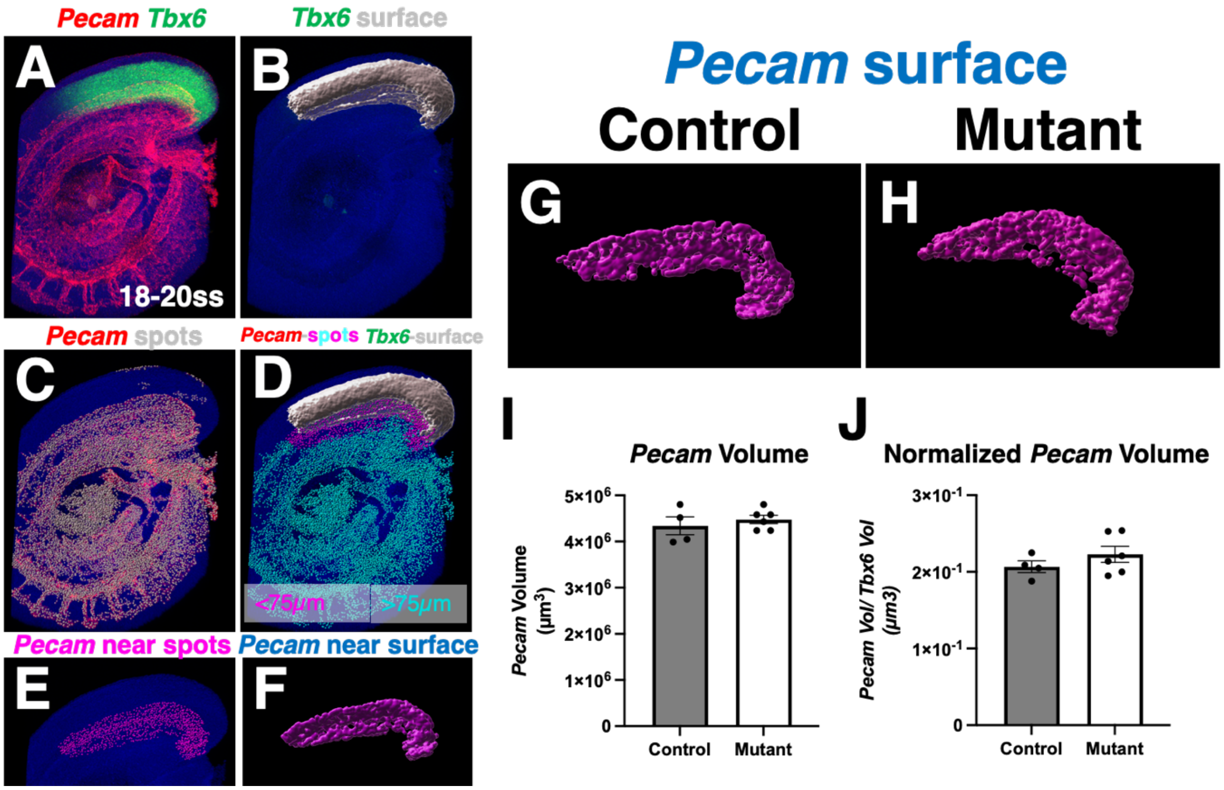


**Fig S2. Mutants have normal vascular development at 18-20ss. (A)** HCR detection of *Pecam* and *Tbx6* mRNA expression at 18-20ss in a control embryo; lateral view of embryo with head removed. **(B)** Tbx6 surface modeling and **(C)** spot detection of *Pecam* expression. **(D)** Detection of *Pecam* positive spots that are within 75μm of *Tbx6* surface (magenta spots **E**) or greater than 75μm from *Tbx6* surface (cyan). **(F)** Surface of *Pecam* vessels near *Tbx6*. **(G,H)** *Tbx6* near *Pecam* surfaces for control and mutant; lateral views, posterior to the right. **(I)** Volume of raw *Pecam* surface or **(J)** *Pecam* surface volume normalized to volume of *Tbx6*; no significant change between control and mutant.

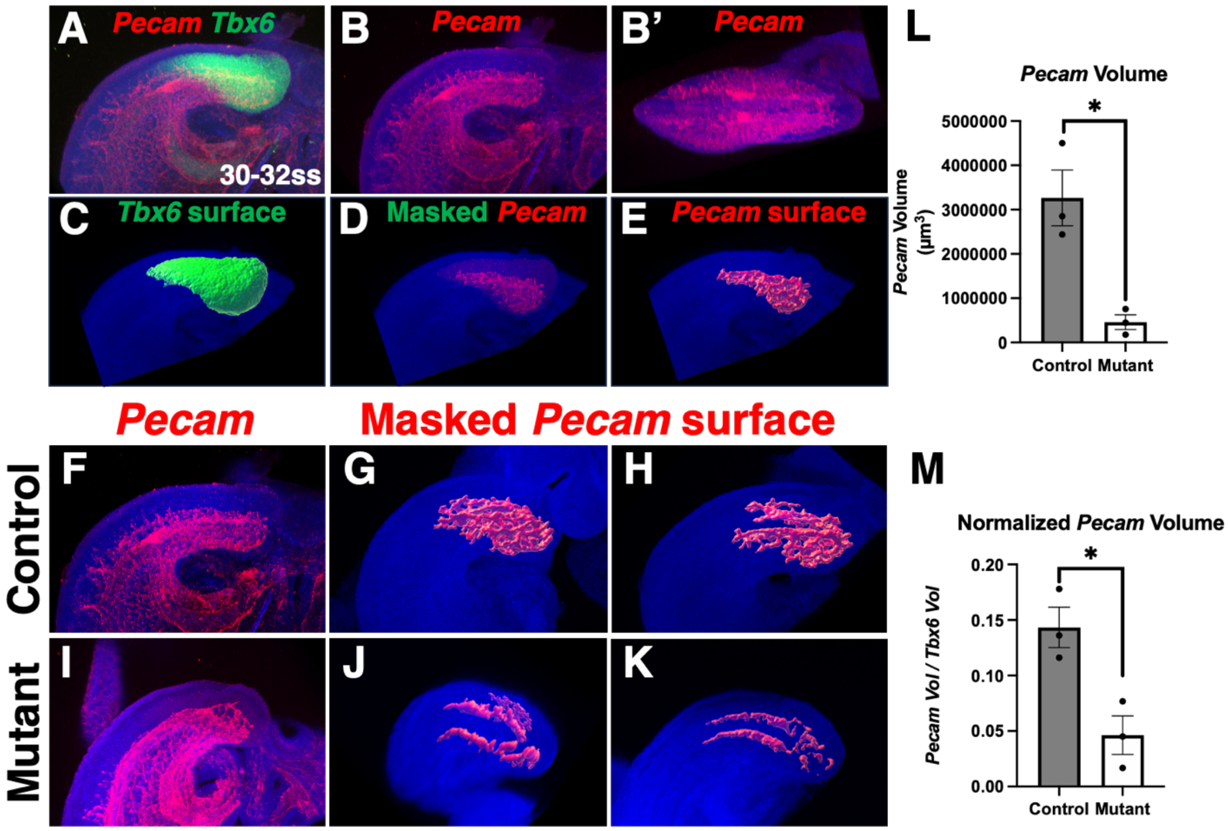


**Fig S3. Mutants have significantly less vascularization of the PSM at 30-32ss. (A)** HCR detection of *Pecam* and *Tbx6* or **(B,C)** *Pecam* only mRNA expression at 30-32ss in a control embryo; (A,B) lateral view or (B') dorsal view of PSM. (C) *Tbx6* surface was used to **(D)** mask *Pecam* expression, and **(E)** surface of *Tbx6* masked *Pecam*; lateral views. **(F,I)** HCR detection of *Pecam* in 30-32ss control and mutant and **(G-K)** PSM masked *Pecam* surfaces from control and mutant; (F,I) lateral and (G,K) dorsolateral views. **(L)** Quantification of *Pecam* volume within PSM and **(M)** *Pecam* volume with PSM normalized to PSM volume; error bars represent SEM, *: $p < 0.05$.

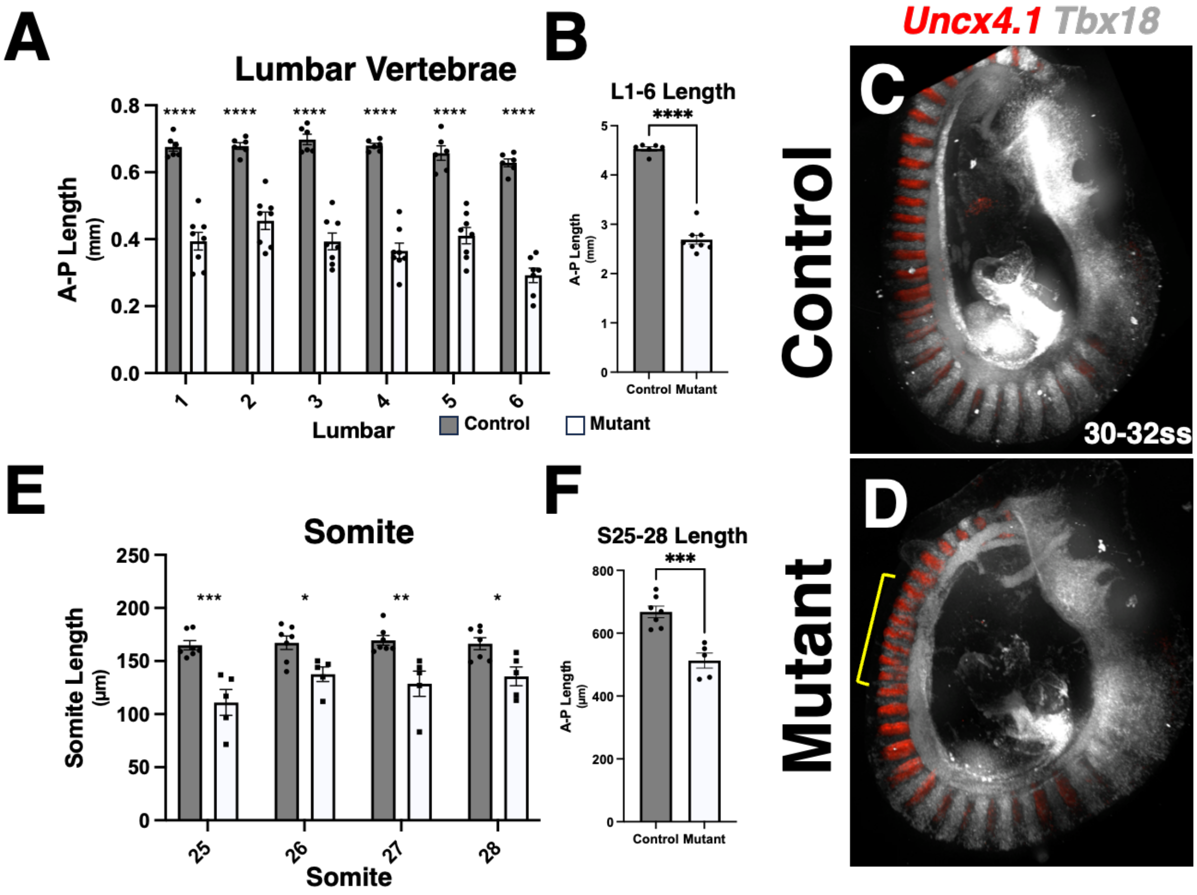


**Fig S4. Mutants have reduced lumbar and somite A-P length. (A)** Individual measurement of A-P length of the first through sixth lumbar vertebrae in E18.5 control (gray bars) and mutant (white bars) E18.5 skeletons or **(B)** sum of lumbar vertebrae length; error bars represent SEM, **** < 0.0001. **(C,D)** HCR detection of *Uncx4.1* and *Tbx18* mRNA expression at 30-32ss in control (n= 7) and mutant (n= 5); bracket highlights region of abnormal somites. **(E)** Individual measurement of A-P length of somite 25 through 28 in 30-32ss control (gray bars) and mutant (white bars) or **(F)** sum of the A-P length of these somites; error bars represent SEM, *** < 0.001.

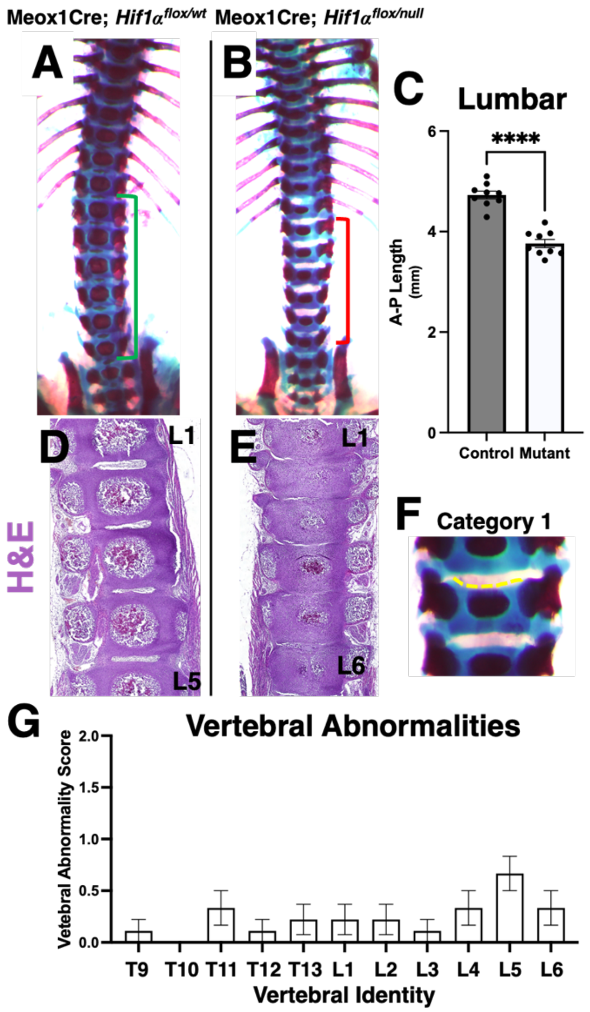


**Fig S5. Meox1Cre; *Hif1α* mutants have reduced vertebral A-P length and mild morphological defects. (A, B)** Skeletons of E18.5 Meox1Cre; *Hif1α* heterozygotes and mutants; brackets indicate lumbar vertebrae. **(C)** Measurement of A-P length of the first through sixth lumbar vertebrae in E18.5 control (gray bar) and mutant (white bar) skeletons; error bars represent SEM, **** < 0.0001. **(D, E)** H & E staining of sections of lumbar vertebrae in E18.5 control and mutant. **(F)** Example of Category 1 vertebral defect; dotted yellow line indicates abnormal shape of articular surface. **(G)** Quantification of vertebral abnormalities based on scoring system (F) from thoracic vertebra 9 (T9) through lumbar vertebra 6 (L6); Meox1Cre; $Hif1\alpha^{flox/null}$ n = 9, error bars represent SEM.

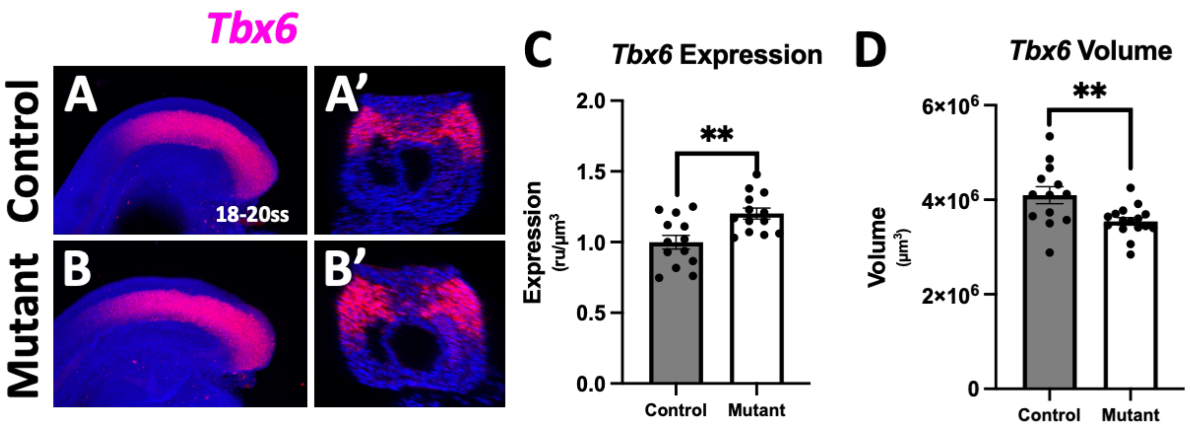


**Fig S6. Mutants have reduced *Tbx6* volume at 18-20ss. (A, B)** Max intensity projection of *Tbx6* mRNA expression at 18-20ss in control and mutant and **(A’, B’)** transverse section through PSM. **(C)** Quantification of *Tbx6* expression within the volume of the *Tbx6* surface and **(D)** volume of *Tbx6* expression domain; error bars represent SEM, **: $p < 0.01$.

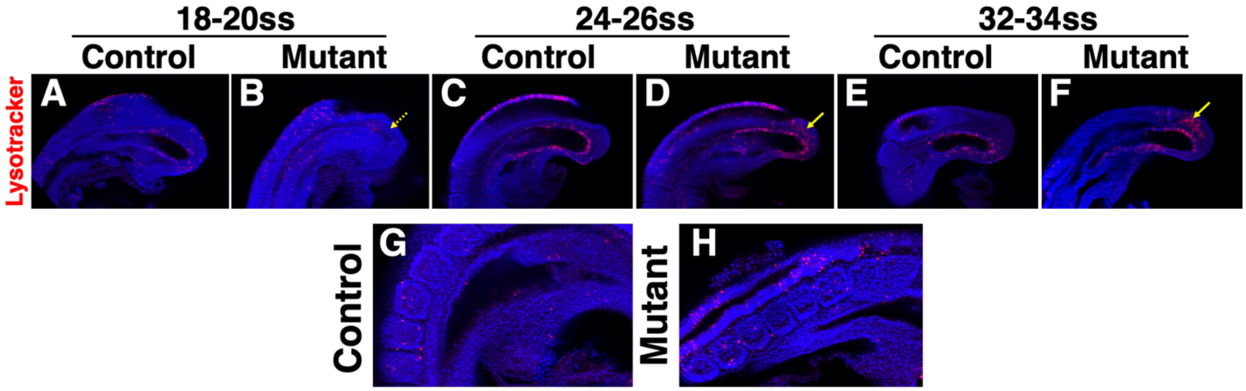


**Fig S7. Mutants have increased cell death in PSM at 32-34ss. (A-F)** Midline sagittal optical sections of Lysotracker stained controls and mutants at indicated stages; yellow arrows indicate region of cell death at 32-34ss, anterior to the left, posterior to the right, 18-20ss: control n= 5, mutant n= 4, 24-26ss: control n= 3, mutant n= 7, 32-34ss control n= 5, mutant n= 3. **(G, H)** Parasagittal optical section of mutant and control 28-30ss embryos showing somites; anterior to the left, posterior to the right, control n= 3, mutant n= 3.

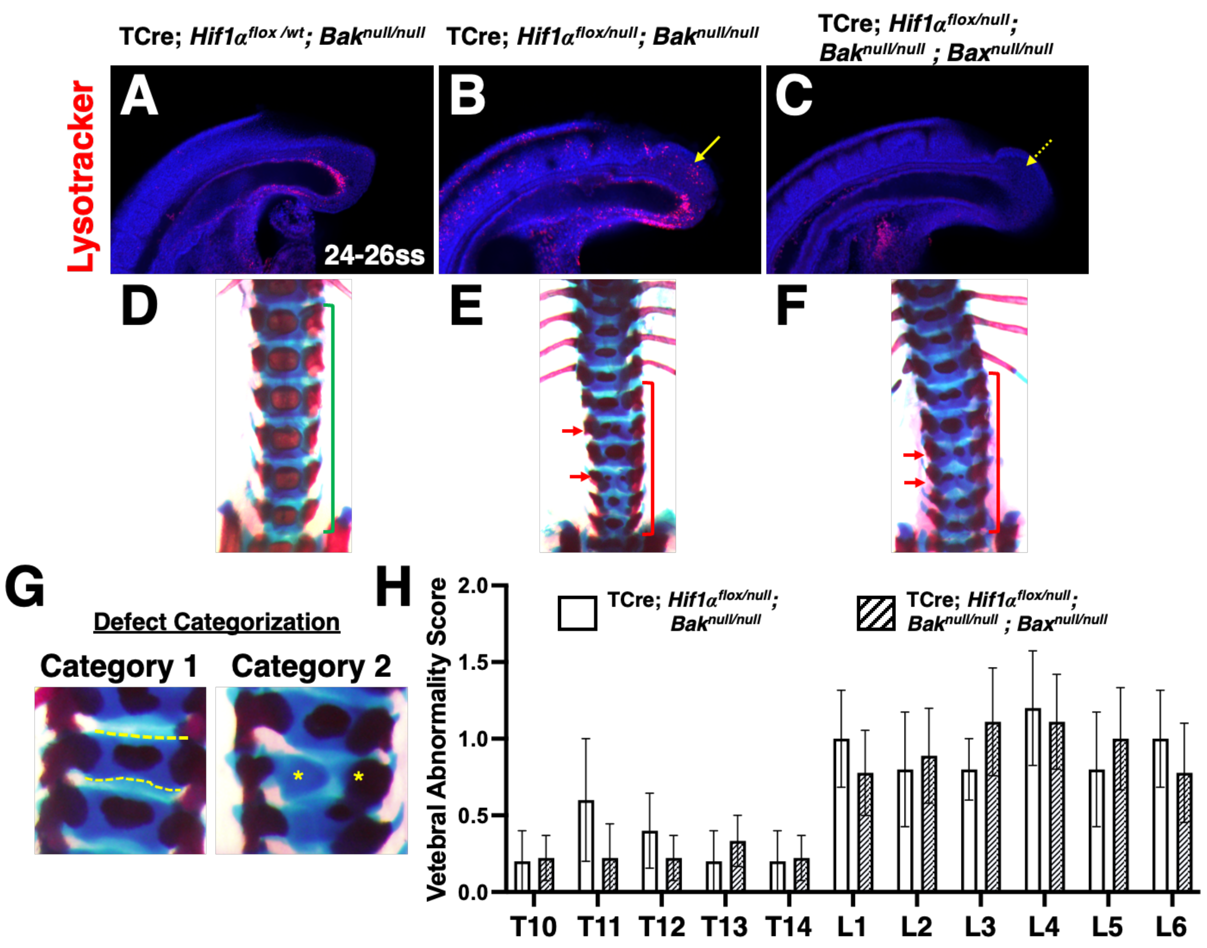


**Fig S8. Removal of *Bak* and *Bax* rescues mutant cell death but does not rescue vertebral morphology. (A-C)** Midline sagittal optical sections of Lysotracker stained 24-26ss embryos with the indicated genotypes; yellow arrows indicate region of cell death in mutants, lateral views, posterior to the right, TCre; *Hif1α*$^{flox/wt}$; *Bak*$^{null/null}$ n= 4, TCre; *Hif1α*$^{flox/null}$; *Bak*$^{null/null}$ n= 3, TCre; *Hif1α*$^{flox/null}$; *Bak*$^{null/null}$ ; *Bax*$^{null/null}$ n= 4. **(D-F)** Skeletal preparations of E18.5 embryos of indicated genotypes showing lumbar region; brackets indicating lumbar region, red arrows indicate highly abnormal vertebrae. **(G)** Examples of Category 1 and Category 2 vertebral defects; dotted yellow lines indicate abnormal shape of articular surfaces, yellow asterisks indicate hemivertebral elements. **(H)** Quantification of vertebral abnormalities based on scoring system, G, from thoracic vertebra 10 (T10) through lumbar vertebra 6 (L6); TCre; $Hif1\alpha^{flox/null}$; $Bak^{null/null}$ n = 5, TCre; $Hif1\alpha^{flox/null}$; $Bak^{null/null}$ ; $Bax^{null/null}$ n = 9, error bars represent SEM, no significant difference was found between genotypes for any vertebral element.

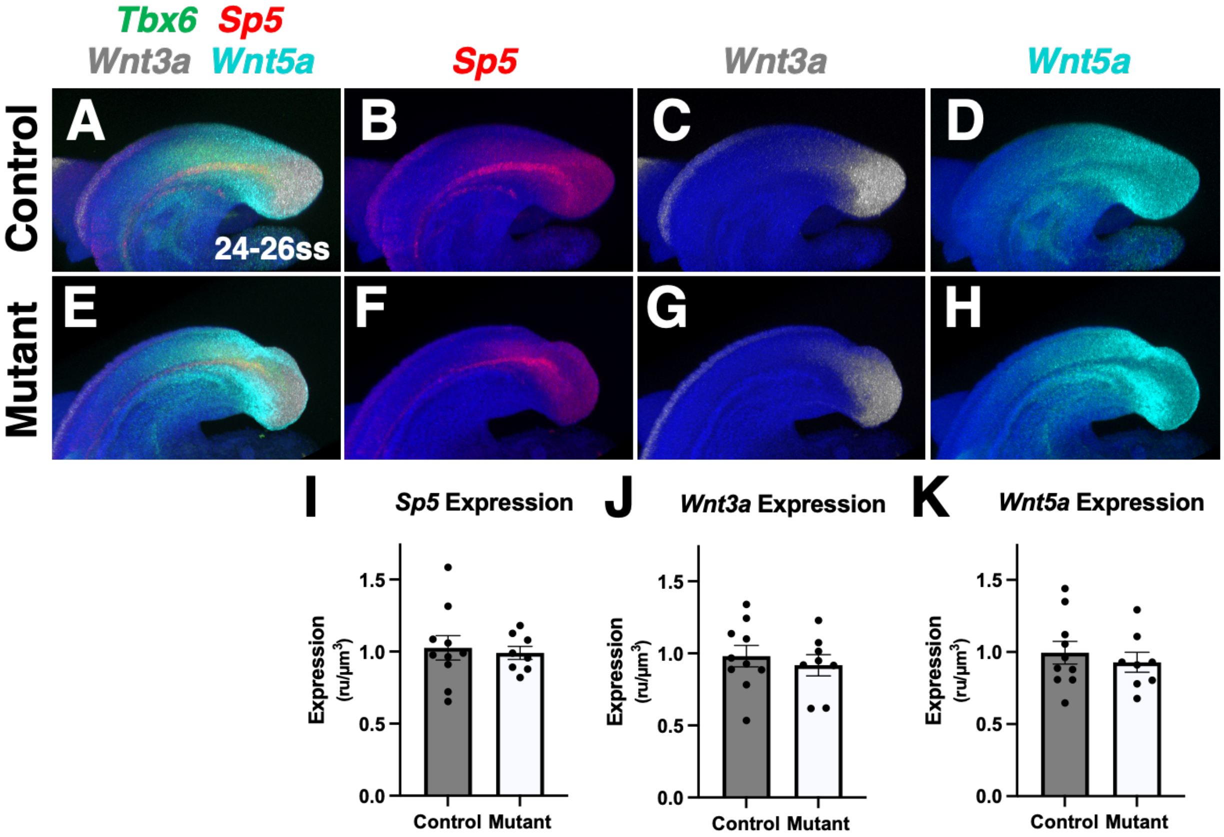


**Fig S9. Wnt signaling is not affected in mutants. (A-H)** Max intensity projections of *Tbx6, Sp5, Wnt3a, and Wnt5a* mRNA expression at 24-26ss in control and mutant; lateral views of PSM, posterior to the right. **(I-K)** Quantification of *Sp5*, *Wnt3a*, and *Wnt5a* mRNA expression within the PSM as defined by *Tbx6* expression; error bars represent SEM, no significant difference was found between control and mutant for any gene expression.

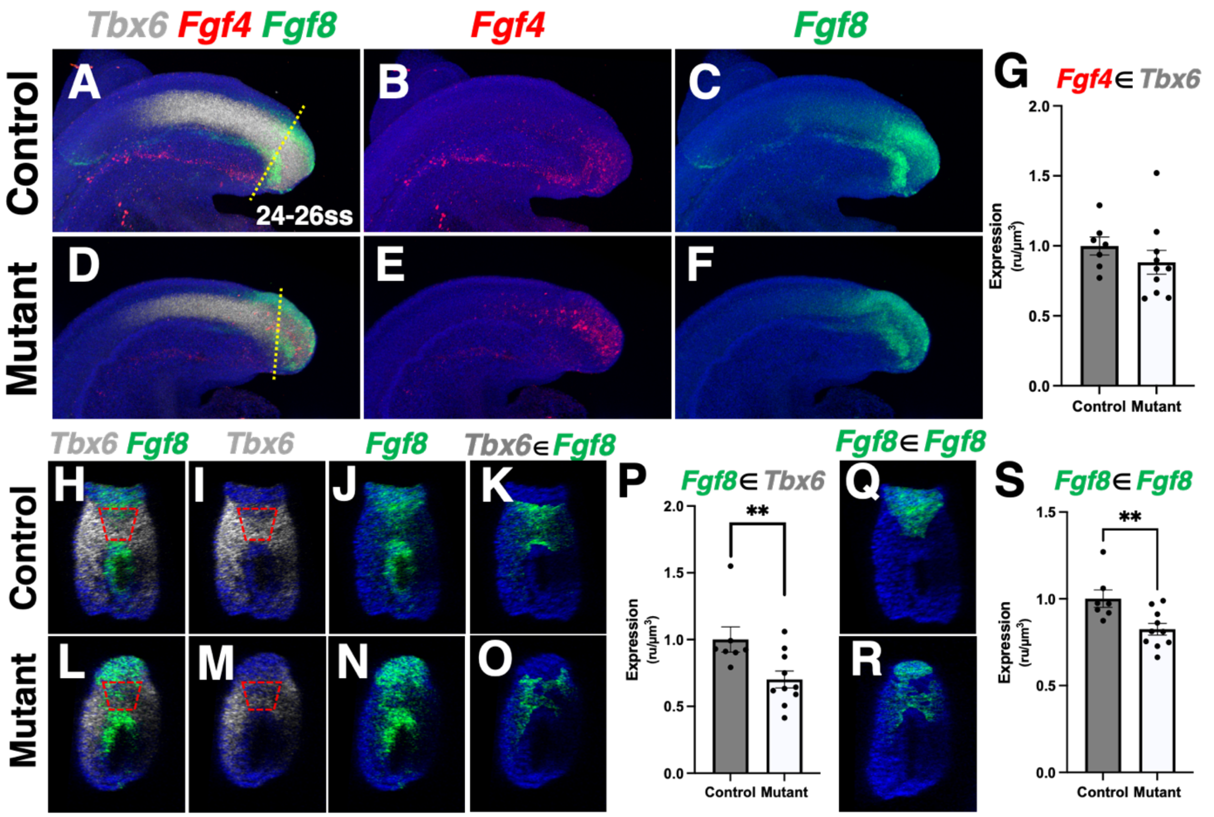


**Fig S10. *Fgf8* but not *Fgf4* expression is reduced in mutants. (A-F)** Max intensity projection of *Tbx6, Fgf4*, and *Fgf8* mRNA expression at 24-26ss in control and mutant; lateral views of PSM, posterior to the right. **(G)** Quantification of *Fgf4* expression within (∈) the PSM as defined by *Tbx6*; error bars represent SEM, no significant difference was found between control and mutant. **(H-J, L-N)** Transverse optical sections at positions indicated by yellow dotted lines in A and D, showing expression of *Tbx6* and *Fgf8*, and **(K, O)** expression within (∈) the *Tbx6* expression domain; red boxes indicate primitive streak location. **(P)** Quantification of *Fgf8* expression within (∈) the PSM as defined by *Tbx6;* **: $p < 0.01$, error bars represent SEM. **(Q, R)** Transverse optical section showing only expression of *Fgf8* within (∈)  its own expression domain (gut tube expression was removed based on morphology). **(S)** Quantification of *Fgf8* expression within its own expression domain; error bars represent SEM , **: $p < 0.01$.

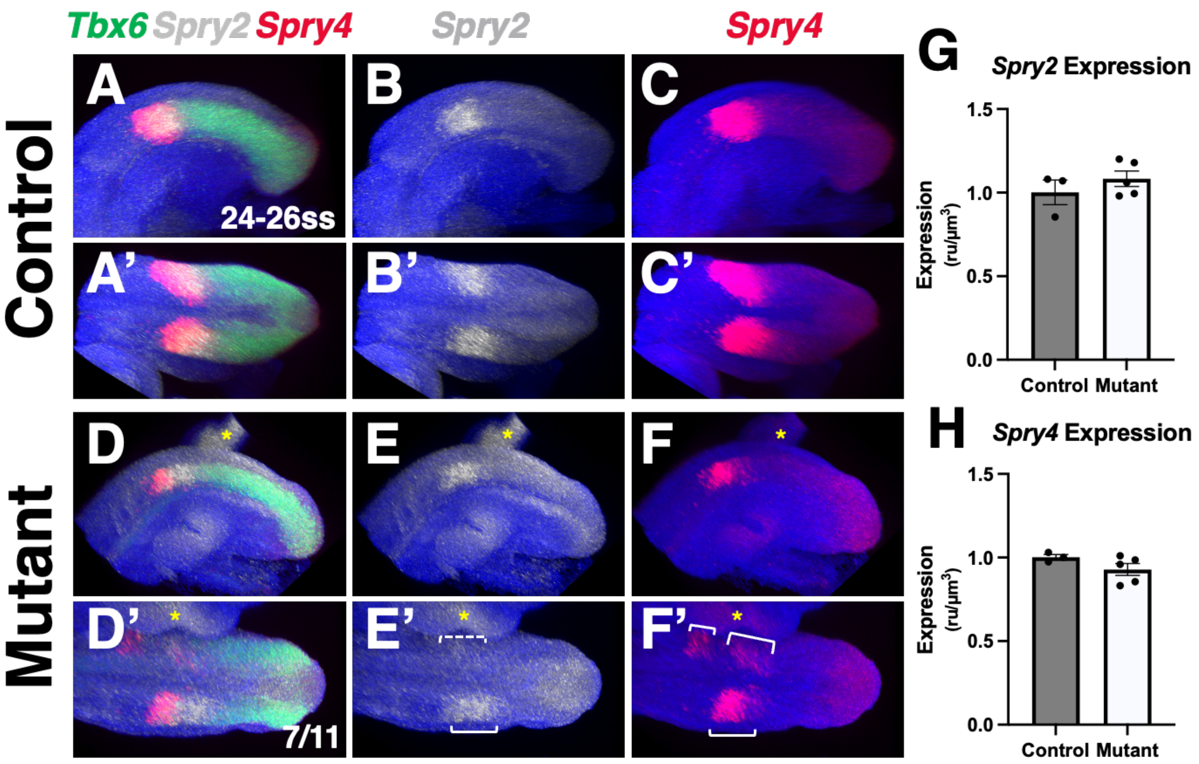


**Fig S11.** ***Fgf*** **responsive gene pattern but not expression is abnormal in mutants.** **(A-F)** Lateral and **(A'-F')** dorsal views of max intensity projections of *Tbx6, Spry2, and Spry4* mRNA expression at 24-26ss in control and mutant; asterisk indicates a piece of the anterior embryo near tail, brackets indicate asymmetric expression domains. **(G, H)** Quantification of *Spry2* and *Spry4* expression within the PSM as defined by *Tbx6*; error bars represent SEM, no significant difference was found between control and mutant.

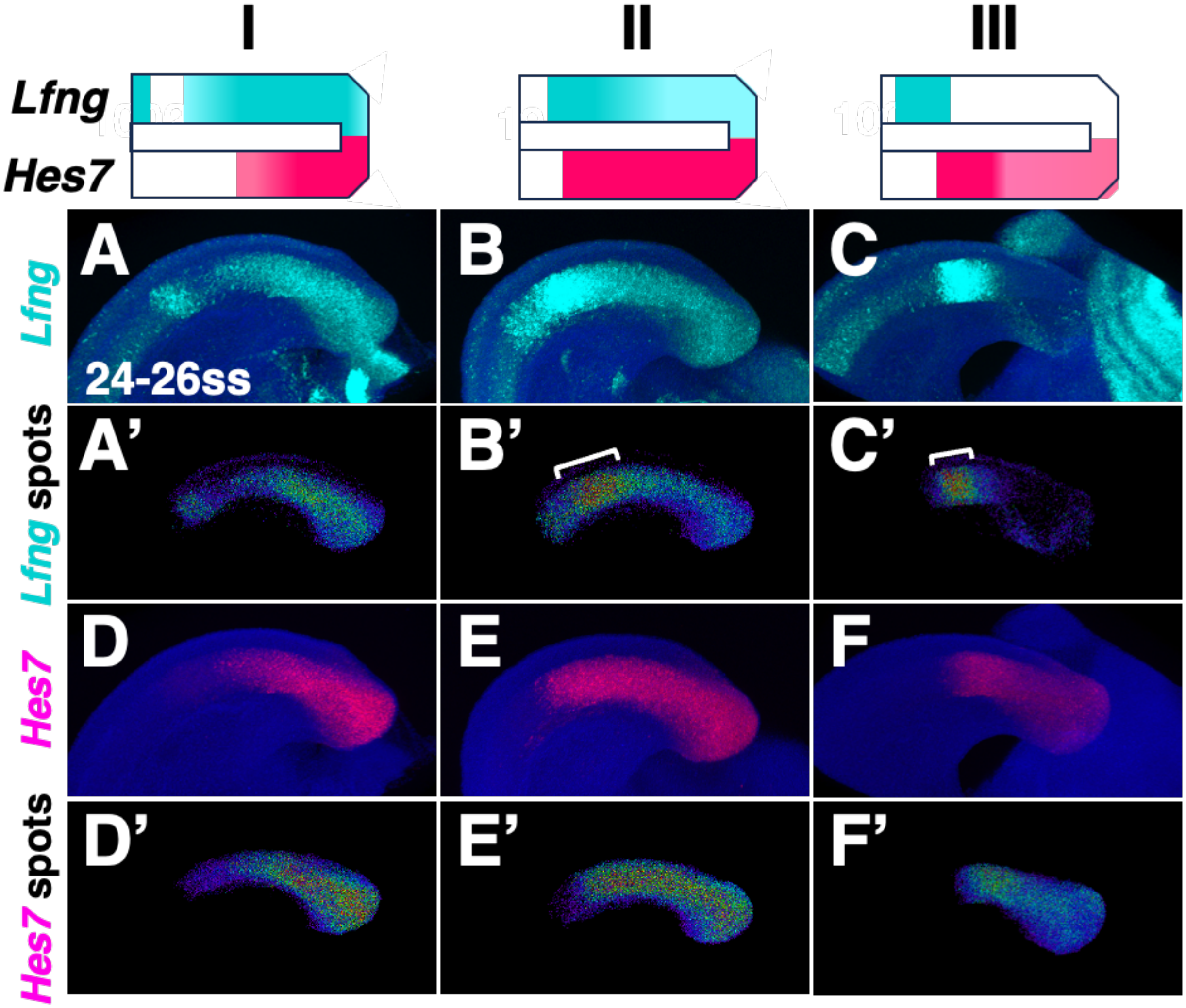


**Fig S12. *Hes7* and *Lfng* oscillation phases. (A-C)** Whole mount HCR detection and **(A’-C’)** spot heatmaps of *Lfng* mRNA expression at 24-26ss in control; brackets indicating higher anterior expression, lateral views of PSM, posterior to the right. **(D-F)** Whole mount HCR detection and **(D’-F’)** spot heatmaps of *Hes7* mRNA expression in same embryos shown in A-C; lateral views of PSM, posterior to the right, n=7. Cartoon of combined phases at top.

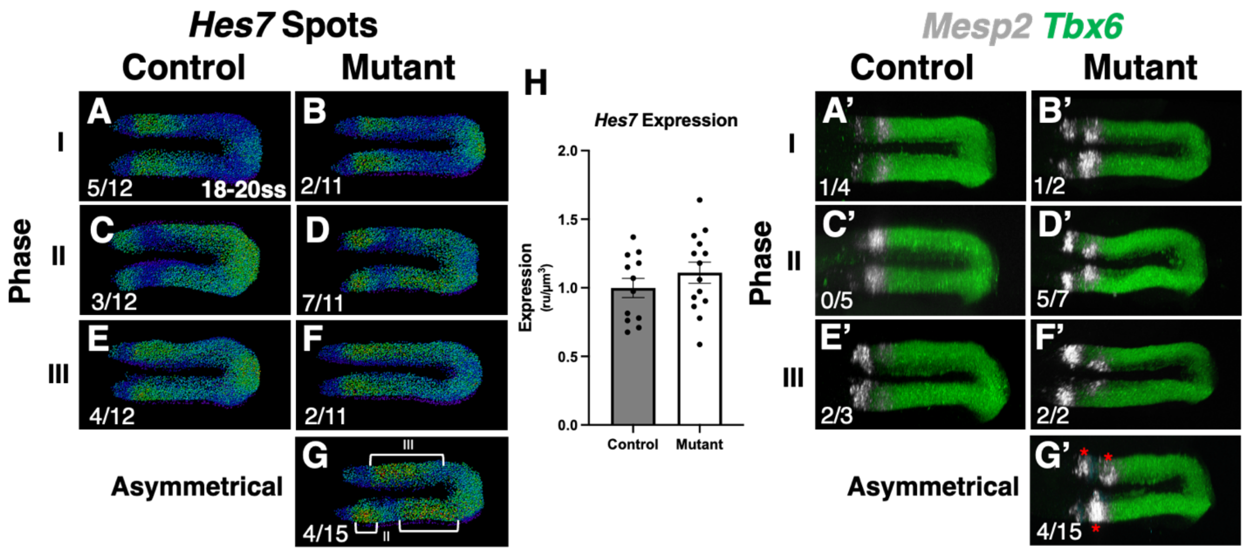


**Fig S13. Mutant *Hes7* and *Mesp2* expression is abnormal at 18-20ss. (A-G)** Spot heatmaps of *Hes7* mRNA expression at 18-20ss in control and mutant sorted by phase; numbers indicate distribution of embryos per phase, note embryos with asymmetric expression were not included in phase distribution numbers in A-F, brackets in G highlight different contralateral phases of expression, dorsal views, posterior to the right. **(H)** Quantification of *Hes7* expression within the PSM as defined by *Tbx6* expression; no significant difference was found between control and mutant, error bars represent SEM. **(A’-G’)** Whole mount HCR detection of *Mesp2 and Tbx6* mRNA expression at 18-20ss in control and mutant sorted by *Hes7* phase; images are same embryos shown in A-G, numbers indicate number of embryos with two *Mesp2* bands per phase, red asterisks indicate asymmetric *Mesp2* bands, dorsal views, posterior to the right.
